# Differential and compensatory roles for type I phosphatidylinositol-4-phosphate-5-kinase isoforms in retinal function and health

**DOI:** 10.64898/2026.05.21.726920

**Authors:** Feng He, Ye Long, Ralph M. Nichols, Samuel M. Wu, Theodore G. Wensel

**Author notes:** **Correspondence:** Theodore G. Wensel, Baylor College of Medicine, One Baylor Plz., Houston, TX 77030.

## Abstract

Phosphatidylinositol (4,5) bisphosphate (PI(4,5)P_2_) plays important roles in development, signaling, intracellular trafficking and regulation throughout the nervous system. Using selective and combined gene ablation strategies, we have determined the roles of this lipid and the kinase isoforms of the PIP5KI family primarily responsible for its synthesis in mouse retina. In rod cells, PI(4,5)P_2_ localizes predominantly to the plasma membrane of inner and outer segments and is enriched in membranes near the synaptic termini. Disruption of the gene encoding the γ PIP5KI isoform, *Pip5k1c*, throughout the developing retina, using Cre expression driven by a Six3 transcription factor-dependent promoter, yields dramatic, but not complete, loss of the protein, with no apparent effects on morphology or function through the first 3-4 months after birth. Slowly progressing photoreceptor degeneration is observed at later ages. Complete loss of the γ isoform in rods, driven by the rhodopsin promoter-based iCre75 transgene, leads to no obvious developmental defects, but results in an earlier-onset rod degeneration. Germ-line ablation of neither the *Pip5k1a* nor the *Pip5k1b* gene leads to any observable morphological defects. Homozygous *Pip5k1a* ablation leads to functional defects in photoreceptors as revealed by reduced a-wave and b-wave amplitudes in the electroretinograms. On the background of rod-specific *Pip5k1c* ablation, *Pip5k1a* deficiency greatly accelerates retinal degeneration. These results reveal a complex interplay among PIP5KI isoforms in ensuring proper photoreceptor function and health, with apparent partial redundancy in fulfilling their critical functions. They underscore the important role of PI(4,5)P_2_ in neuronal signaling and homeostasis.

**Significance Statement:** Phosphatidylinositol(4,5)P_2_, PI(4,5)P_2_, plays essential roles in nervous system development and function, but its roles in retina have been unknown. This study combines biochemistry, mouse genetics, light- and electron microscopy to reveal both specific and redundant functions for PIP_2_ formed by different kinase isoforms in the mammalian retina. It has implications for retinal function, disease and therapy, and for the broader field of phosphoinositide regulation.

## Introduction

Phosphatidylinositol-(4,5)-bisphosphate, abbreviated here as PI(4,5)P_2_, is a low-abundance lipid that plays multiple functional roles in cells, including direct modulation of activity of ion channels, receptors and other membrane proteins, re-organization of the actin-based cytoskeleton, and coordination of membrane traffic (Di Paolo and De Camilli, 2006). The latter activity ranges from initiation of endocytosis or exocytosis to intracellular vesicle transport, to ciliary membrane organization, to fusion of vesicles with lysosomes prior to degradation and/or recycling of their contents. It also serves as the substrate for the generation of second messengers: inositol-(1,4,5)-trisphosphate (InsP_3_) and diacylglycerol upon the receptor-dependent activation of phospholipase C, and phosphatidylinositol-(3,4,5)-trisphosphate (PIP_3_) upon the receptor-dependent activation of PI-3-kinase. These and other functions are mediated through specific interactions of PI(4,5)P_2_ or clusters of PI(4,5)P_2_ with hundreds of different proteins containing PI(4,5)P_2_-binding domains(Itoh and Takenawa, 2002; Sasaki et al., 2007; Monteiro et al., 2014; Hammond and Balla, 2015; Martin, 2015; Lystad and Simonsen, 2016; Chandra and Collins, 2019; Eitzen et al., 2019; Feng et al., 2019; Tsuji et al., 2019; Overduin et al., 2022).

In the brain, PI(4,5)P_2_ has been shown to be important for proper regulation of exocytic vesicle release at synapses (Di Paolo et al., 2004) and, in dorsal root ganglion neurons, it has been implicated in growth cone morphology (Yamazaki et al., 2013). In the retina, its function has not been thoroughly explored. However, it is known that certain enzymes involved in regulation of PI(4,5)P_2,_ such as the PI-phosphatases INPPE (Bielas et al., 2009; Sharif et al., 2021) and synaptojanin (Van Epps et al., 2004; Holzhausen et al., 2009) are essential for proper retinal structure, function and health, as are the PI(4,5)P_2_-binding proteins Tubby and TUPLP1 (North et al., 1997; Hagstrom et al., 1998; Hagstrom et al., 1999). INPP5E and Tubby family members (Mukhopadhyay and Jackson, 2011) have been implicated in cilium function. The G proteins G_αq_ and Gα11 and the PI(4,5)P_2_-directed phospholipase C isozymes downstream of them have been reported to be present in the retina (Peng et al., 1997), but their functional roles in outer retina are not well understood. In intrinsically photosensitive ganglion cells, this pathway plays an important role in melanopsin signaling (Xue et al., 2011; Jiang et al., 2018).

The bulk of cellular PI(4,5)P_2_ a arises from the action of isoforms of the PIP5KI family, which catalyze the phosphorylation of PI(4)P at the 5-position on the inositol ring (Doughman et al., 2003; Volpicelli-Daley et al., 2010). There are three genes encoding these enzymes in both mice and humans, with the nomenclature switched between the two species with respect to α *vs.* β isoforms, *i.e.*, mouse *Pip5k1b*, *Pip5k1a* and *Pip5k1c* are the orthologs of human *PIP5K1A*, *PIP5K1B* and *PIP5K1C*, respectively, and encode, respectively, PIP5KIβ, PIP5KIα, and 1PIP5KIγ in mouse. There are splice variants of these isoforms that have previously been reported to have differential spatial distributions and functions (Heck et al., 2007).

Previous results with knockouts and knockdowns have suggested that the different PIP5KI-encoding genes have different but partly redundant functions (Wang et al., 2007). In mice *Pip5k1c* knockouts are lethal at embryonic or perinatal stages (Wang et al., 2007; Wang et al., 2008a), whereas *Pip5k1b* and *Pip5k1a* knockouts are viable (Sasaki et al., 2005; Wang et al., 2007; Wang et al., 2008a). However, *Pip5k1a* knockouts display sperm defects with reduced fertility, and *Pip5k1b* plus *Pip5k1a* double-knockouts have even lower fertility (Hasegawa et al., 2012).

## Results

### Phototransduction-dependent enhancement of PI(4,5)P_2_ levels by light

We extracted acidic phospholipids from rod outer segments (ROS) isolated from either dark-adapted or light-adapted mice and assayed the extracts for levels of PI(4,5)P_2_ (Fig. 1A). The assay used was similar to one published previously for PI(3)P, but used a construct based on the high affinity PI(4,5)P_2_ -binding pleckstrin homology (PH) domain of phospholipase-Cδ (PLCδ); see Methods for details. The results of such experiments consistently revealed significantly higher levels of PI(4,5)P_2_ in light-adapted, as compared to dark-adapted ROS. The same assays carried out in light-adapted retinas revealed that this light-induced increase in PI(4,5)P_2_ levels was greatly reduced in phototransduction-deficient mice with both copies of the gene for the phototransduction G protein, transducin inactivated, *i.e.*, *Gnat1*^-/-^ mice (Fig. 1B). Thus, light exposure triggers significant increases in steady-state levels of PI(4,5)P_2_ through a mechanism substantially driven by the same phototransduction cascade that elicits physiological light responses of rod cells. Experiments measuring levels of PI(4)P in parallel with PI(4,5)P_2_ levels, indicated that its levels are increased more moderately in the light, and that PI(4,5)P_2_ levels are comparable to those of PI(4)P in the light, but are much lower in the dark (Supplementary Fig. S1).

**Figure 1.**
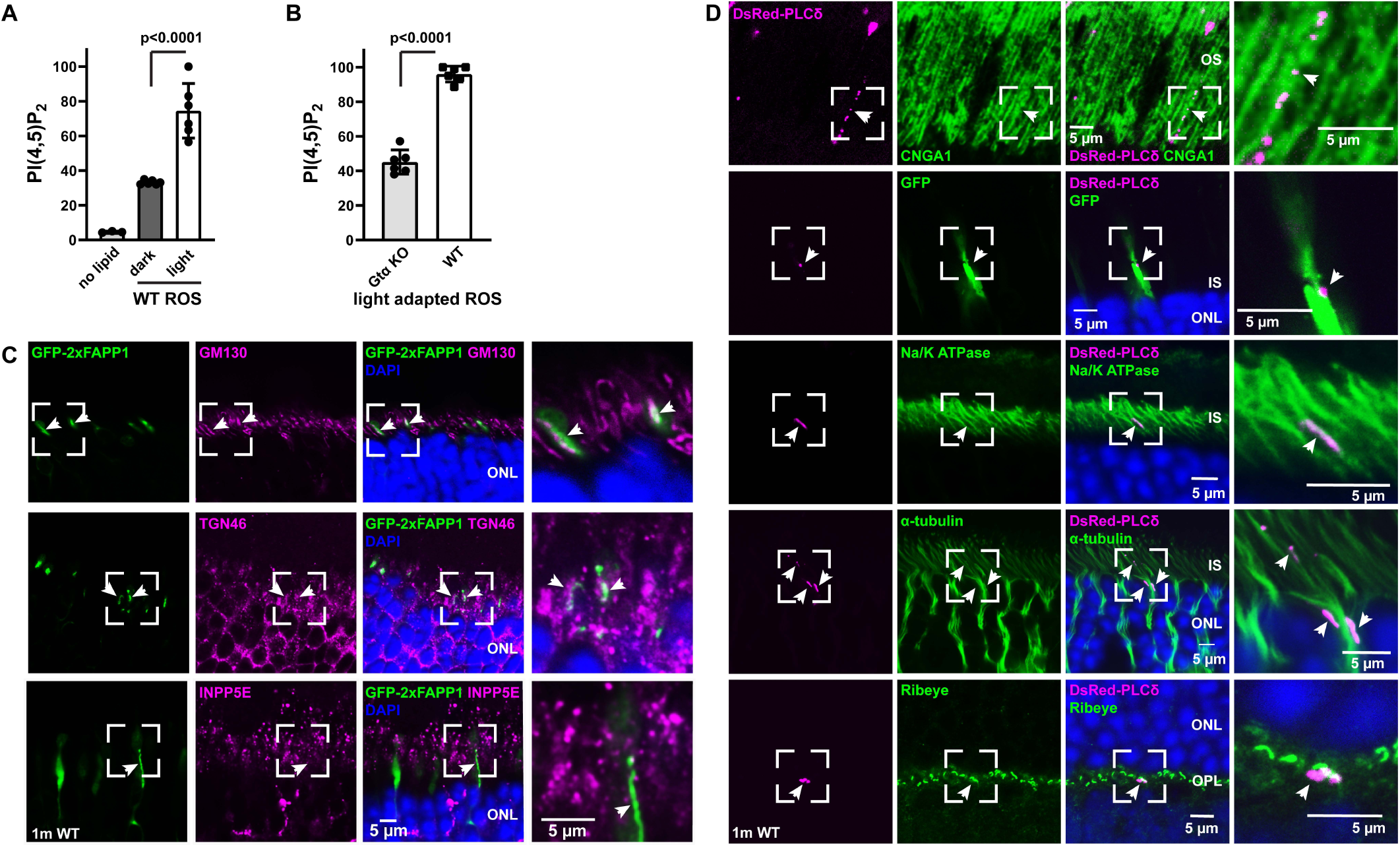
Light regulation and localization of PI(4)P and PI(4,5)P_2_ in mouse rod cells. **(A, B)** Light regulation of PI(4,5)P_2_ in mouse retina. PI(4,5)P_2_ level in purified ROS was determined by phosphoinositide ELISA (*n* = 6). (**A)** ROS from WT mice, (**B**) ROOS from *Gnat^-/-^* mice (transducin KO). (**C, D**). Four-phosphate-adaptor protein 1 (FAPP1) that binds to PI(4)P, and phospholipase C-delta (PLCδ) pleckstrin homology domain that binds specifically to PI(4,5)P_2_ were used to visualize PI(4)P and PI(4,5)P_2_ in mouse photoreceptor cells. Probe expression was directed by a plasmid encoding a fluorescent protein-tagged phosphoinositide binding domain, eitherFP-FAPP1 or DsRed-PLCδ, under control of a mutant (A7T) proximal opsin promoter, which induces probe expression at <10% of the level induced by the WT promoter. Plasmids were introduced by sub-retinal injection and electroporation of P0 WT CD1 mice (selected due to absence of pigment). In retinas probed with the PLCδ-derived probe, a construct encoding GFP was co-injected to facilitate identification of successfully transfected cells (“GFP” in second row of panel **D**). Mouse retina sections from 1-month-old injected mice were immunostained with indicated antibodies.

### Localization in outer retina of PI(4,5)P_2_, PIPI-kinases, and their substrate, PI(4)P

In order to determine the localization of PI(4,5)P_2_ and its precursor PI(4)P in the retina we electroporated plasmids encoding fusions of fluorescent proteins with phosphoinositide-binding domains specific for PI(4)P or PI(4,5)P_2_ into the retinas of neonatal mice (P0,or 0 days PN) following microinjections, using methods described previously (Matsuda and Cepko, 2004; He et al., 2016; He et al., 2019; Agosto and Wensel, 2021; He et al., 2025). For PI(4)P we used a tandem repeat of four-phosphate-adaptor protein 1 (FAPP1) fused to enhanced Green Fluorescent Protein (GFP-2xFAPP1), and for PI(4,5)P_2_, we used the phospholipase C-delta (PLCδ) PH domain fused to dsRed (dsRed- PLCδ).

Initial experiments using a chicken β actin promoter with a CMV enhancer (“CAG” promoter(Matsuda and Cepko, 2004)) or a minimal rhodopsin promoter (Pawlyk et al., 2005; Lee et al., 2010) to drive expression led to excessive over-expression of the probe, with the result that diffuse cytoplasmic signal masked any specific membrane-bound signal. We therefore adopted a modified version of this promoter that, based on previous findings (Lee et al., 2010), was expected to yield only moderate expression, and observed highly specific membrane labeling in rods (Fig. 1C, D).

As in other cell types, strong signal for PI(4)P was associated with the Golgi apparatus and trans-Golgi network (Fig. 1C), accompanied by weaker signal associated with plasma membrane and other unidentified intracellular membranes. Interestingly, although there is extensive overlap between PI(4)P signal and the Golgi marker GM130, there were regions *adjacent* to the GM130 signal, but not containing it, that showed strong signal for PI(4)P. Signal for the trans-Golgi network marker, TGN46, also overlapped only partially, but was consistently found adjacent to PI(4)P signal. PI(4)P signal did not overlap with that for the phosphoinositide phosphatase INPP5E.

In contrast, signal for PI(4,5)P_2_ in the inner and outer segments was exclusively observed in the plasma membrane, where it strongly co-localized with the plasma membrane-resident cyclic nucleotide-gated channel in the outer segment, or with the plasma membrane-restricted Na/K ATPase in the inner segment (Fig. 1D). Surprisingly, PI(4,5)P_2_ signal was not observed uniformly throughout the plasma membranes of transfected rods but rather was restricted instead to extensive but localized patches of membrane. Although evidence has been reported of PI(4,5)P_2_ clustering has been reported (reviewed by (Wen et al., 2021)), including a recent report in cerebellar neurons (Eguchi et al., 2023), it is usually distributed fairly uniformly in plasma membranes, and the striking localization we observe is not observed in other cell types. We cannot rule out the possibility that this strong clustering is an artifact of some kind, possibly due to dsRed aggregation or fixation. However, the evidence for PI(4,5)P_2_ presence in both the inner and outer segment membranes and its exclusion from internal membranes seems robust. Its presence in only a very small fraction of total photoreceptor membranes is consistent with its low levels in rod cells, as compared to other cell types (Wensel, 2020).

As the Type I PIP kinases have been proposed to be responsible for the bulk of PI(4,5)P_2_ synthesis in most cell types (Doughman et al., 2003; Volpicelli-Daley et al., 2010), we probed mouse retina with antibodies specific for the α and γ isoforms (Fig. 2). We also tried antibodies reported to be specific for the β isoform. Unfortunately, when we compared results between WT and β knockout mice, these β antibodies were found to be specific in immunoblots (see below), but not in immunofluorescence experiments. Therefore, retinal immunostaining with these antibodies is not shown, because evidence indicates it is non-specific.

**Figure 2.**
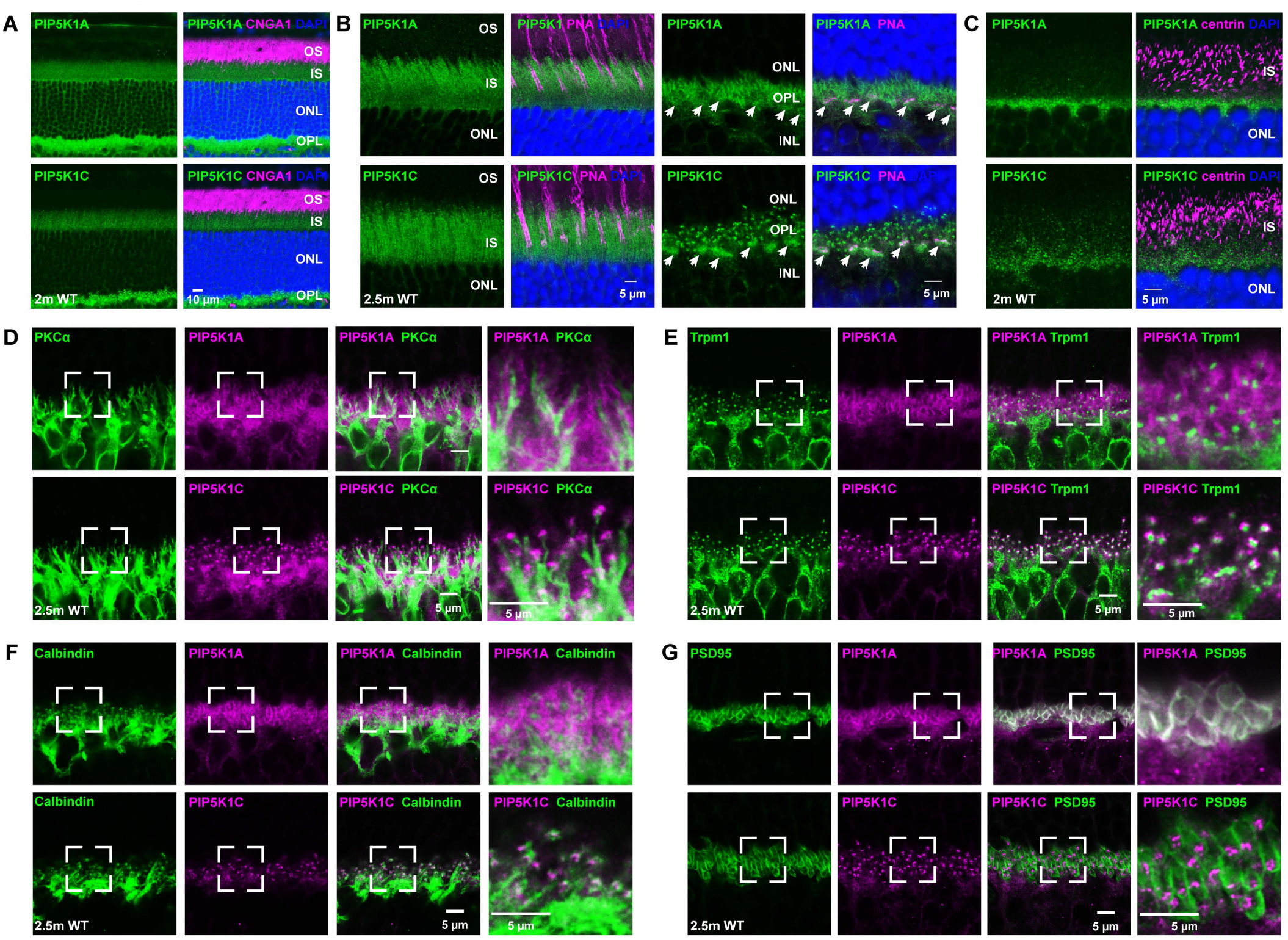
Localization of PIP5K1α and PIP5K1γ in wild-type photoreceptor cells. **(A)** PIP5K1α and PIP5K1γ (labeled PIP5K1A and PIP5K1C), are localized in the inner segment and outer plexiform layer of mouse photoreceptor cells. **(B)** Higher magnification images of the inner segment (left two panels) and outer plexiform layer (right two panels) reveal distinct patterns between PIP5K1α and PIP5K1γ in the synaptic region (OPL). PNA labels cone sheaths. **(C)** PIP5K1γ labels puncta in the inner segment but does not colocalize with connecting cilium marker, centrin. **(D-G)** PIP5K1α localizes to the plasma membrane of the presynaptic terminal, whereas PIP5K1γ is positioned near the synaptic ribbon of the photoreceptor cells and the tips of bipolar cell dendrites. PKCα, was used to mark rod bipolar cell cytoplasm (**D**), TRPM1 as a marker for ON-bipolar cell dendritic tips and endoplasmic reticulum (**E**), calbindin (**F**) as a marker of horizontal cells, and PSD95 as a marker of pre-synaptic photoreceptor terminals. Strong co-localization of PIP5K1α is observed only for PSD95, (**G**). Subsequent labeling of rod-specific KO of *Pip5k1c* (Fig. 3) confirmed OPL PIP5K1γ signal to be predominantly pre-synaptic.

[*Note on nomenclature: Throughout, to enhance legibility of figure labels, “*PIP5K1A,” “PIP5K1B”, and “PIP5K1C*”, are used in labels on figures to refer to the mouse proteins,* PIP5KIα, PIP5KIβ, PIP5KIγ*, respectively or, when followed by “*KO*,” are used to refer to retinas from mice with germ-line (for* α *or* β *isoforms) or conditional (for* γ*) inactivation of the genes encoding them. Samples from mice heterozygous for any of knockouts are indicated in figure labels as “het.” “*PN*” is used throughout to indicate “post-natal” age*]

Signal for the α isoform of PIP5KI (Fig. 2) was seen across multiple retinal layers. No signal above background was reliably detected in the outer segment layer (OS), but there was clearly faint signal in the inner segment and inner nuclear layers (Fig. 2A). The IS staining appeared to be present throughout the cell but was concentrated in the IS plasma membranes. Signal was particularly bright in the outer plexiform layer (OPL). Inner segment staining appeared to include both rods and cones, marked by peanut agglutinin (PNA), but there was no co-localization with staining by centrin antibodies, a marker for the connecting cilium (CC) in photoreceptors. The OPL staining (Fig. 2B-G) strongly co-localized with staining for PSD-95, a pre-synaptic marker in the OPL (Fig. 2G) (Koulen et al., 1998). Co-staining for rod ON-bipolar cell marker PKCα (protein kinase Cα, Fig. 2D), cone sheath marker PNA (Fig. 2B) and horizontal cell marker calbindin (Fig.2F) showed PIP5KIα to be largely pre-synaptic. There were no puncta corresponding to the synapse-forming dendritic tips of ON-bipolar cells marked by TRPM1 (Fig. 2E). There was also diffuse cytoplasmic staining, not strongly co-localized with any of these markers, in the OPL.

PIP5KIγ signal (Fig. 2) was also found at low levels in the IS and ONL, but was absent in the outer segments and CC. The signal was clearly enhanced in the IS plasma membranes (Fig. 2 and Supplementary Fig. 2A). There were somewhat larger numbers of small puncta in the IS than observed for PIP5KIα, but these were similar to those observed in controls (*i.e.*, possibly due to non-specific background) and almost none were associated with the CC marked by centrin staining. PIP5KIγ signal was more intense in the OPL and inner plexiform layer (IPL) than in the IS. In the OPL (Fig. 2B-G), PIP5KIγ signal strongly labeled synaptic regions, including both rod spherules, identified by their pattern of doublet punctae, and the cone pedicles, marked by PNA and characteristic morphology. There was also fainter and diffuse labeling of interneuron cell bodies (Fig 2D-G). PIP5KIγ labelling did not overlap PSD-95 signal (Fig. 2G) but was closely associated with TRPM1 labeling of ON-bipolar cell dendritic tips (Fig. 2E). Additional experiments (see below) revealed this TRPM1-associated label to be presynaptic.

### Single PIP5K1 knockouts: Germline and conditional knockouts of genes encoding PIP5KI isoforms

In order to determine the roles of the PIP5KI isoforms and the pools of PI(4,5)P_2_ they generate in the mammalian retina, we studied mice with germ-line inactivation of P*ip1ka and Pip1kb*, and pan-retinal or rod-specific knockout of *Pip1kc*. While knockouts of *Pip5k1a* and *Pip5k1b* genes are viable through adulthood, (Sasaki et al., 2005; Wang et al., 2008a) germ-line knockouts of Pip5k1c are not (Di Paolo et al., 2004; Wang et al., 2008b). Therefore, we obtained mice bearing a “floxed” allele of *Pip5k1c* (*i.e.*, one containing loxP sites flanking essential regions) and used *Cre* transgenes to drive conditional knockouts. Pan-retinal inactivation of *Pip1kc* was attempted using a Cre transgene driven by the regulatory sequences from the *Six3* transcription factor gene, which has been used for this purpose previously (Burns et al., 2008; Zhang et al., 2008; Fuhrmann et al., 2009; Zhang et al., 2015; Syc-Mazurek et al., 2017; Dilan et al., 2019; Cao et al., 2021; Huang et al., 2022; Yan et al., 2022; Bright et al., 2024). Although the *Six3* gene is known to be expressed by retinal precursor cells early in retinal development and to drive transcription of its target genes at that stage (Furuta et al., 2000; Peng et al., 2017; Duan et al., 2018), our results as described below suggest that PIP1Kγ protein is not completely eliminated in rod cells until well after terminal differentiation of rods, which are the last retinal neurons to reach that step (Cepko, 1996; Morrow et al., 1998). We achieved effective elimination of the γ isoform in rods using the iCre75 transgene (Li et al., 2005), which is fully activated and achieves efficient *Pip1kc* inactivation as soon as rhodopsin expression is activated around post-natal day 11, yielding effectively quantitative excision of floxed alleles by post-natal day 18.

Immunoblots (Fig. 3A) at 2 months PN confirmed the effectiveness of the α and β isoform knockouts at eliminating those proteins. Immunoblots also confirmed a large reduction in the amount of the γ isoform in the retina of the pan-retinal Six3-Cre-driven knockout, However, the reduction in total retinal PIP5KIγ protein in the iCre75-driven knockout was too low to be detected. Immunofluorescence using a Cre-specific antibody confirmed high levels of Cre in the outer nuclear layer, which is predominantly made up of rod nuclei, in the iCre75 transgenics. In the Six3-driven knockouts, high levels of Cre expression were observed in the inner nuclear layer and ganglion cell layer, but not in the outer nuclear layer at age 2 months PN (Fig. 3B).

**Figure 3.**
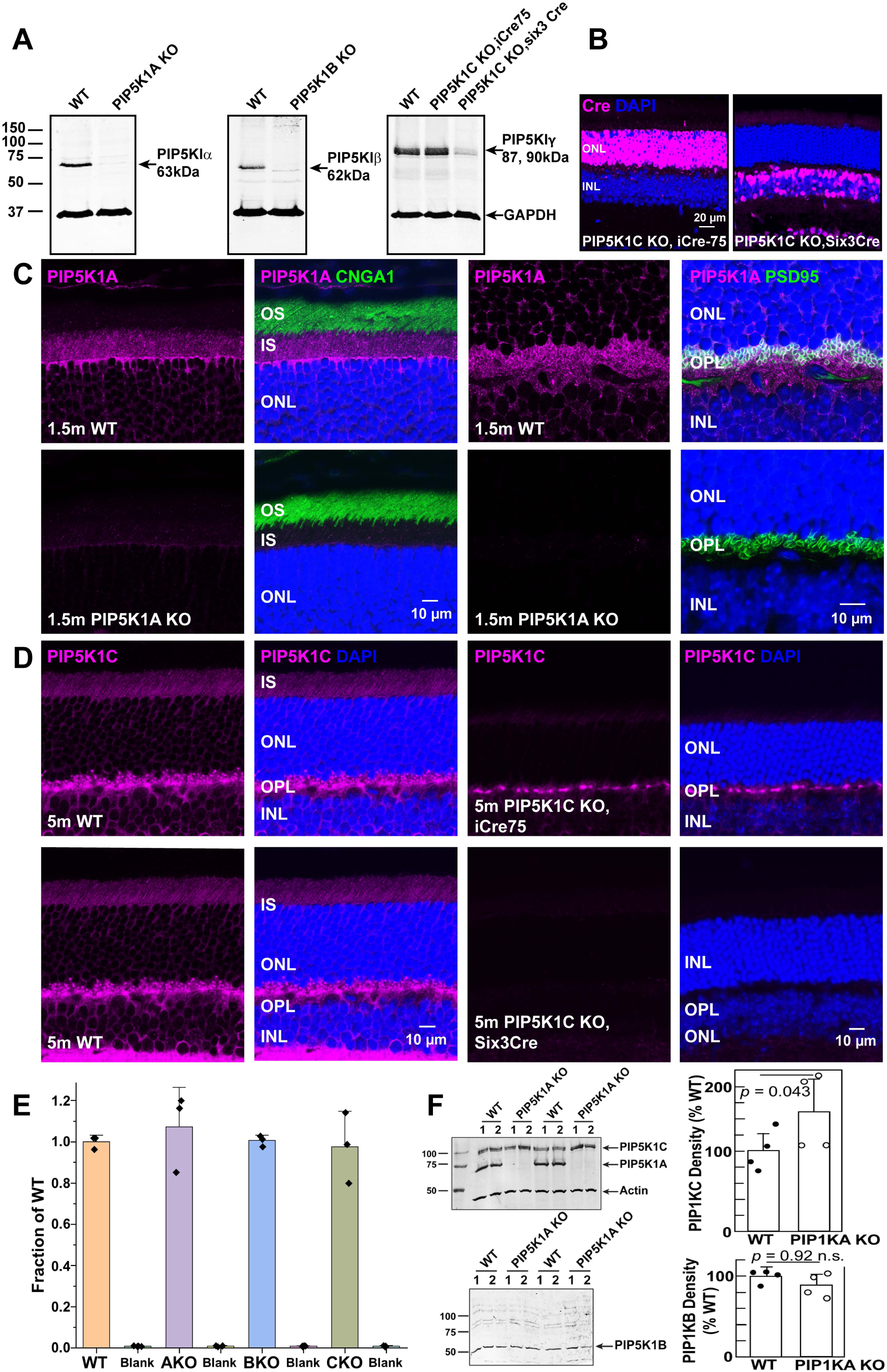
Depletion of PIP51 kinase isoforms in single-isoform knockouts of PIP5K1α, PIP5K1β, and in retina-specific PIP5K1γ deletion. **(A)** Immunoblot analysis of whole mouse retina lysates comparing wild-type and single-isoform PIP5K1 knockouts shows protein levels for PIP5K1α, PIP5K1β, and PIP5K1γ at 2 months PN. GAPDH serves as a loading control. **(B)** Immunofluorescence for Cre recombinase at 6 months PN using iCre75 for photoreceptor-specific (left) and Six3-Cre for pan-retinal knockout of PIP5K1γ (right, 2 months PN). **(C)** Immunofluorescence images of photoreceptor layer (left two columns) or OPL (right two columns) for WT and PIP5K1α knockout mice. Loss of PIP5K1α immunoreactivity was observed in the inner segment and outer plexiform layer of PIP5K1α knockout retina sections. These results also serve to validate the specificity of immunostaining with the PIP5K1α antibody. Cross-reactivity in immunostaining observed for available antibodies for PIP5K1β precluded similar experiments for knockout of the PIP5K1β isoform. **(D)** Immunofluorescence images of IS, ONL, OPL and INL for WT (left two columns) and rod-specific PIP5K1γ KO (right two columns) driven by iCre75 or Six3Cre. PIP5K1γ was absent in photoreceptor cells in iCre75-driven knockout (upper two panes), and its expression is lost in the entire retina in Six3-Cre-driven knockout at 5 m PN (lower two panels). These results also serve to validate the specificity of immunostaining with the PIP5K1γ antibody and to confirm that the OPL PIP5K1γ is predominantly pre-synaptic. **(E)** Assays of total retinal PI(4,5)P_2_ levels in single-isoform knockouts do not show significant depletion. *AKO*, ELISA results from retinas lacking PIP5K1α; *BKO*, ELISA results from retinas lacking PIP5K1β; *CKO*, ELISA results from retinas lacking PIP5K1γ in rods. Error bars indicate ±s.d. **(F)** Immunoblot results for PIP5K1γ and PIP5K1β levels in PIP5K1β in PIP5K1α KO retinas. For each condition, *n* = 4; bar heights represent mean values and error bars the s.d.

Immunofluorescence experiments at 1.5 months PN confirmed the absence of detectable PIP5KIα in the retinas of α knockout mice (Fig. 3C). At 5 months of age, no PIP5KIγ could be observed in the retinas of Six3-Cre-driven γ knockout mice by immunofluorescence (Fig. 3D). At 2 months the Six3-Cre-driven PIP5KIγ knockout mice had readily detectable PIP5KIγ signal in both the inner segments and the outer plexiform layer, and at 3 months, they still had faint PIP5KIγ signal in the inner segments (Supplementary Fig.2A, B). These results indicate that the Six3-Cre construct used did not lead to elimination of PIP5KIγ protein until well into adulthood.

At 5 months (Fig. 3D), retinas from the iCre75-driven knockouts were devoid of detectable PIP5KIγ protein in the photoreceptor inner segments or in the outer nuclear layer. PIP5KIγ staining was also lost in the region of the outer plexiform layer immediately adjacent to the ONL, although some remained in the more inner regions of the OPL, consistent with loss of presynaptic PIP5KIγ protein in rods, and with its continued presence in interneurons of the OPL. These results confirmed both the effectiveness of the knockouts and the specificity of the antibodies used. Despite the complete loss of the β isoform signal in immunoblots (the faint residual band in Fig. 3A corresponds to the wrong molecular weight), immunofluorescence experiments indicated no loss of signal, leading to the conclusion that the antibodies used are not specific for PIP5Iβ under immunofluorescence staining conditions.

Assays of PI(4,5)P_2_ levels in retinas from each of the PIP5KIα or PIP5KIβ single knockouts or the rod-specific PIP5KIγ knockout revealed no significant decreases in overall levels in light-adapted retinas at 4 months PN (Fig. 3E). These results are consistent with compensatory PI(4,5)P_2_ synthesis catalyzed by the remaining isoforms in retinas with each single knockout. Quantitative immunoblotting (Fig. 3F) indicated that there is a modest increase (36%) in PIP5KIγ levels in PIP5KIα knockout retinas, but, if anything a slight but non-significant *decrease* in PIP5KIβ levels. Further studies will be needed to determine whether there is compensatory upregulation of activity for any isoforms in the single knockouts, or possibly altered localization of any isoform in response to elimination of another. A limitation of these results is that our measurements would likely not be sensitive to localized changes in PI(4,5)P_2_

### Morphological effects of single knockouts

The germ-line knockouts of PIP5KIα and PIP5Iβ had no apparent morphological defects through six months of age (Fig. 4A, B). In contrast, the iCre75-driven knockouts of PIP5KIγ, while morphologically normal at 2 months of age, displayed significant loss of photoreceptor nuclei at 4 months, and reduction by about half by 6 months after birth (Fig.4C). The Six3-Cre-driven knockouts of PIP5KIγ did not show obvious signs of degeneration at 4 months but displayed significant shortening of rod outer segments and some loss (∼1-3 rows) of photoreceptor nuclei at 6 months (Fig. 4F). Thus, knockout of PIP5KIγ leads to slow degeneration of photoreceptor cells (Fig.4D), but not to any obvious morphological defects in other retinal layers. Single knockout of either PIP5KIα or PIP5KIβ leads to no apparent degeneration of any part of the retina at any age studied. No obvious developmental defects were observed in any of the single knockout lines.

**Figure 4.**
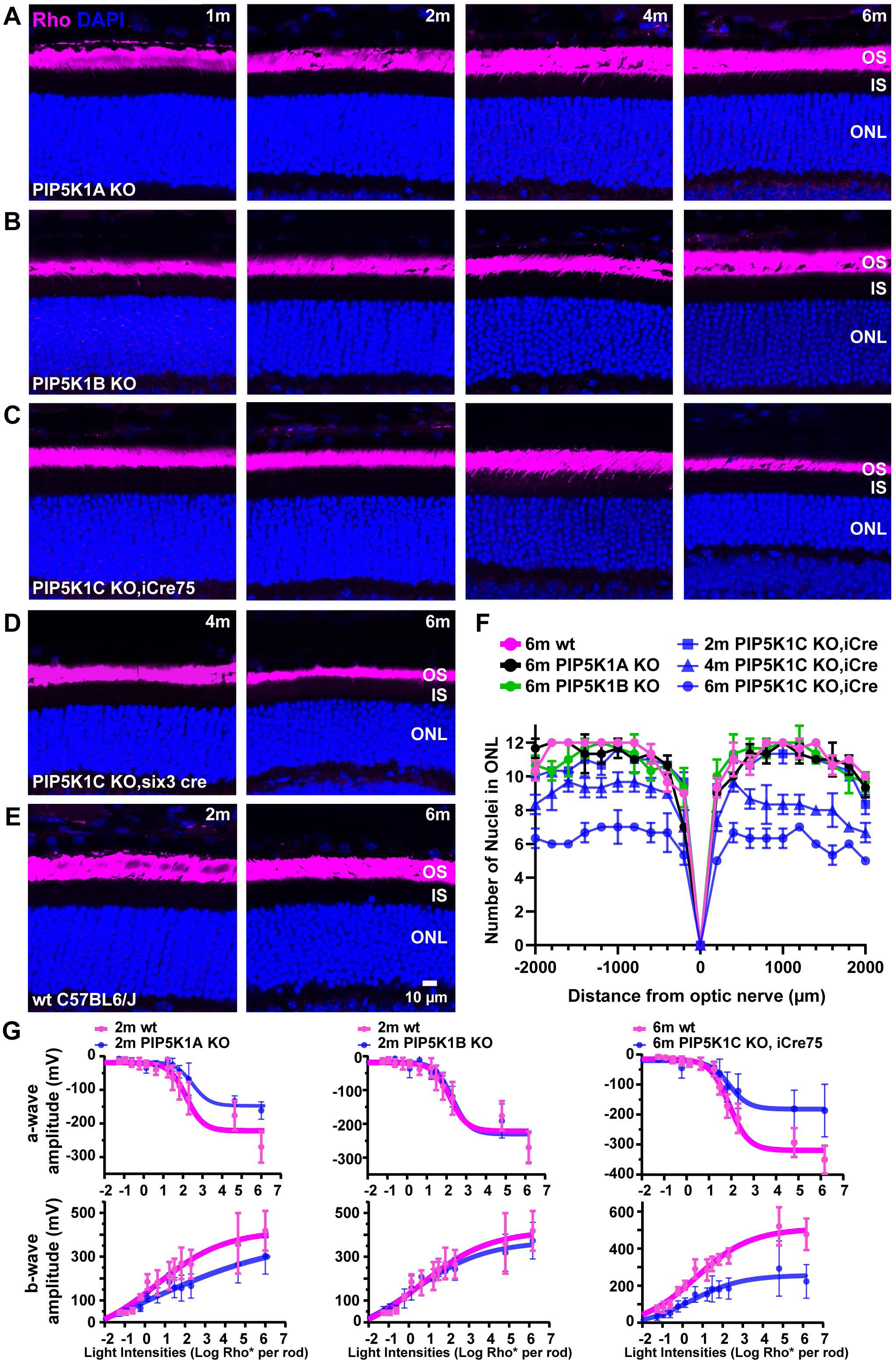
Effects on photoreceptor cells of single isoform PIP5K1α or PIP5K1β knockout, and of retina-specific PIP5K1γ knockout. **(A, B)** Retina sections from PIP5K1α or PIP5K1β knockout mice stained with rhodopsin antibody (Rho) and DAPI showed no significant photoreceptor degeneration up to 6 months of age. **(C)** In photoreceptor-specific (iCre75-driven) PIP5K1γ knockout mice, 20–30% of photoreceptor cells were lost at 4 months, increasing to 50% by 6 months. **(D)** Retina-specific (Six3-Cre-driven) PIP5K1γ knockout mice exhibited approximately 10–20% photoreceptor cell loss at 6 months. **(E)** Control retina sections stained with Rho and DAPI. **(F)** “Spidergram” plots show outer nuclear layer (ONL) thickness versus distance from the optic nerve at 2, 4, and 6 months PN. Data are presented as mean ± SEM; *n* = 3 mice per age group. **(G)** Electroretinography (ERG) revealed reduction of a-wave and b-wave amplitudes in 2-month-old PIP5K1α KO, and 6-month-old PIP5K1γ KO mice. No significant changes were observed in PIP5K1β knockout mice. Error bars represent mean ± SEM, *n* = 4–6 mice. Scale bar in panel **E** applies to panels **A-E**.

### Functional defects in single knockouts

Electroretinography revealed functional defects for some of the single-knockout genotypes (Fig. 4G). Not surprisingly, the rod-specific PIP5KIγ knockouts, which suffered loss of rods, had diminished ERG amplitudes, as compared to WT animals. There were amplitude decreases in both their a-waves, which reflect rod hyperpolarization, and their b-waves, which largely arise from ON-bipolar cells immediately downstream of rods, under dark adapted conditions. At 6 months of age, when rod numbers were reduced by about half, the a-wave amplitudes were also reduced by about half at the higher intensity ranges, where signal-to-noise ratios allow valid comparisons. The b-wave amplitudes were similarly reduced. No ERG deficits were observed in PIP5KIβ knockouts, which also displayed no loss of photoreceptor cells.

Of great interest was the result that PIP5KIα knockouts did show significantly diminished a-wave and b-wave amplitudes at 2 months after birth (Fig. 4G), despite the absence of any observable morphological defects or cell death. <u>These results suggest</u> <u>that PI(4,5)P_2_ specifically generated by PIP5KIα is important for the efficiency of the</u> <u>light response in rod cells.</u>

Because of the high levels of PIP5KI proteins in the outer plexiform layer, we used immunofluorescence to look for mislocalization of proteins in that region (Figs. 5, 6A). In neither the PIP5KIα knockouts nor the PIP5KIβ knockouts was there any mislocalization of the pre-synaptic markers PSD95 (Koulen et al., 1998) or Bassoon (Brandstätter et al., 1999), the postsynaptic marker, TRPM1 (Agosto et al., 2018b; Agosto et al., 2018a), or the rod-bipolar cell marker, PKCα (Haverkamp and Wässle, 2000; Ruether et al., 2010). In contrast, all these markers showed mislocalization within the outer nuclear layer at 4 or 6 months of age in the iCre75-driven PIP5KIγ knockouts, consistent with the well-documented phenomenon of neuronal remodeling and bipolar cell dendritic “sprouting” resulting from loss of photoreceptors (*e.g.,* (Marc et al., 2007)) as well as in normal aging retina (Liets et al., 2006).

**Figure 5.**
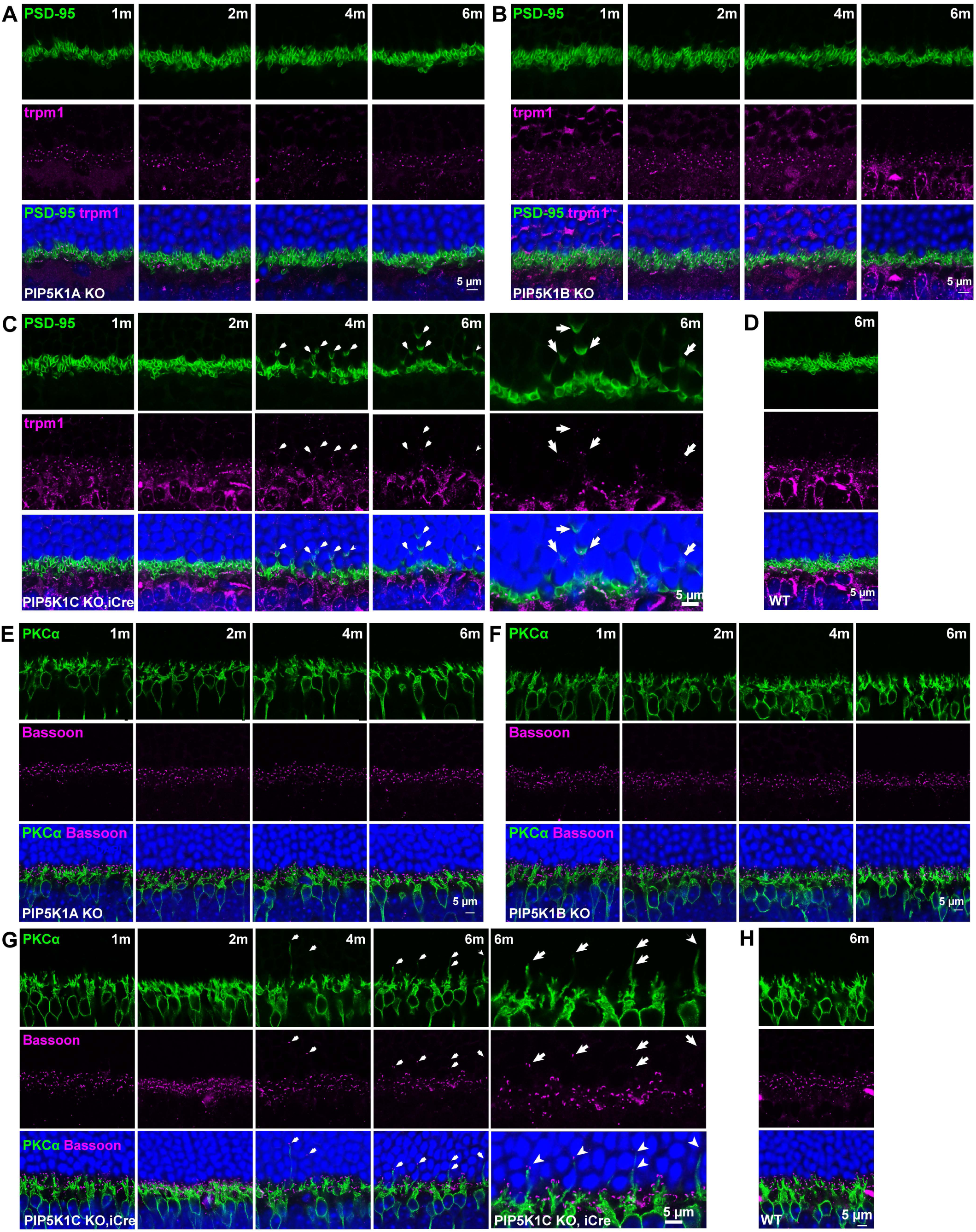
Effects of global (PIP5K1α, PIP5K1β), or photoreceptor-specific PIP5K1γ knockout on the outer plexiform layer (OPL). **(A, B)** Co-immunostaining of photoreceptor presynaptic terminal marker PSD-95 and bipolar cell marker TRPM1 revealed no noticeable morphological changes in the OPL compared to wild-type mice in PIP5K1α and PIP5K1β knockout mice (ages 1–6 months PN; ages are given in the upper right of each series of panels in **A**-**F**). **(C)** In PIP5K1γ knockout mice, photoreceptor axons retracted into the outer nuclear layer (ONL), accompanied by rod bipolar cell sprouting (arrows) at 4–6 months, concomitant with loss of photoreceptor cells. **(E, F)** Rod bipolar cell marker PKCα and photoreceptor synaptic ribbon marker Bassoon were normally expressed in the OPL of PIP5K1α and PIP5K1β knockout mice up to 6 months. **(G)** In iCre-75-driven PIP5K1γ knockout mice, rod bipolar dendrites extended into the ONL, and the photoreceptor cell ribbon marker Bassoon mislocalized to the same region (white arrows), indicating shortened photoreceptor axons in the absence of PIP5K1γ.

**Figure 6.**
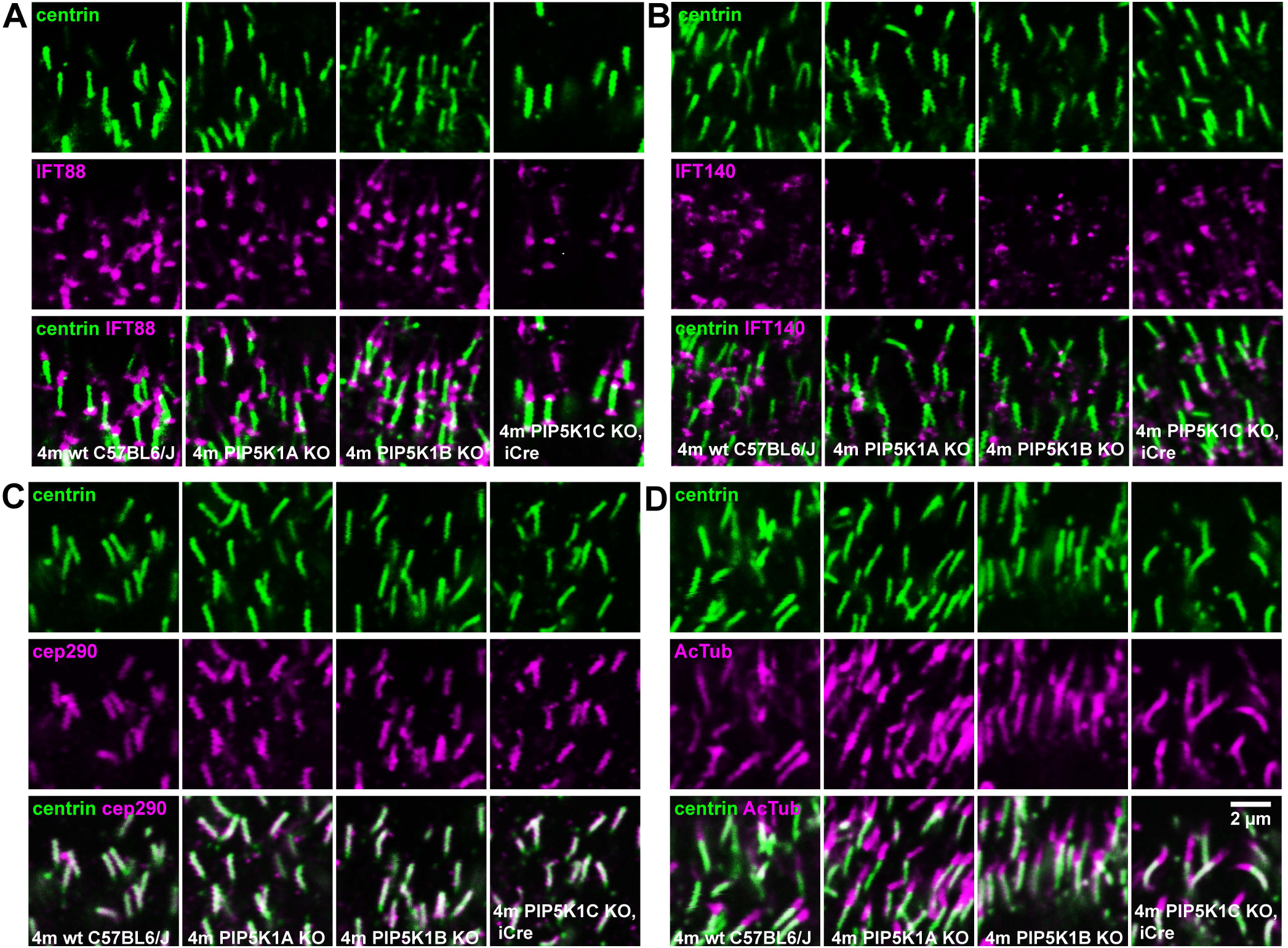
Effects of single knockouts on protein distributions in the photoreceptor connecting cilia. **(A, B)** Intraflagellar transport proteins IFT88 and IFT140 showed normal distributions in the connecting cilia of PIP5K1α, PIP5K1β, and PIP5K1γ knockout mice at 4 months. **(C, D)** Ciliary proteins centrin, CEP290, and acetylated tubulin were similarly distributed in PIP5K1α, PIP5K1β and PIP5K1γ knockouts compared to wild-type mice. Scale bar in lower right panel of **D** applies to **A-C**.

Similar immunofluorescence imaging experiments (Fig. 6B-E) revealed no evidence for protein mislocalization or trafficking defects in the connecting cilium due to single knockouts. Intraflagellar transport proteins IFT88 and IFT140 showed normal distributions in the connecting cilia of all three single knockout mice at 4 months (Fig. 6B, C). Ciliary proteins centrin, CEP290, and acetylated tubulin were also normally distributed in PIP5K1α, PIP5K1β and PIP5K1γ knockouts as compared to WT at 4 m PN (Fig. 6D, E).

### Effects of double knockouts

Effects of double knockouts were studied in mice homozygous for both global *Pip5k1a* KO and *Pip5k1b* KO, or for genotypes consisting of one of these global knockouts and *Pip5k1c* conditional KO driven by either Six3-Cre, or ICre-75 (Figs. 7–9, Supplementary Figs. S3-S7). Combining *Pip5k1b* knockout with either *Pip5k1a* KO or rod-specific *Pip5k1c* KO did not exacerbate the phenotypes beyond what was observed with the *a* or *c* single knockouts (Fig. 7G, I, J, L). Out to 6 months of age there was no apparent loss of nuclei in *Pip5k1b* plus *Pip5k1a* double KO animals (Fig. 7I). Loss of *Pip5k1b* function also did not exacerbate the ERG phenotype of the rod-specific *Pip5k1c* KO (Fig. 7L).

**Figure 7.**
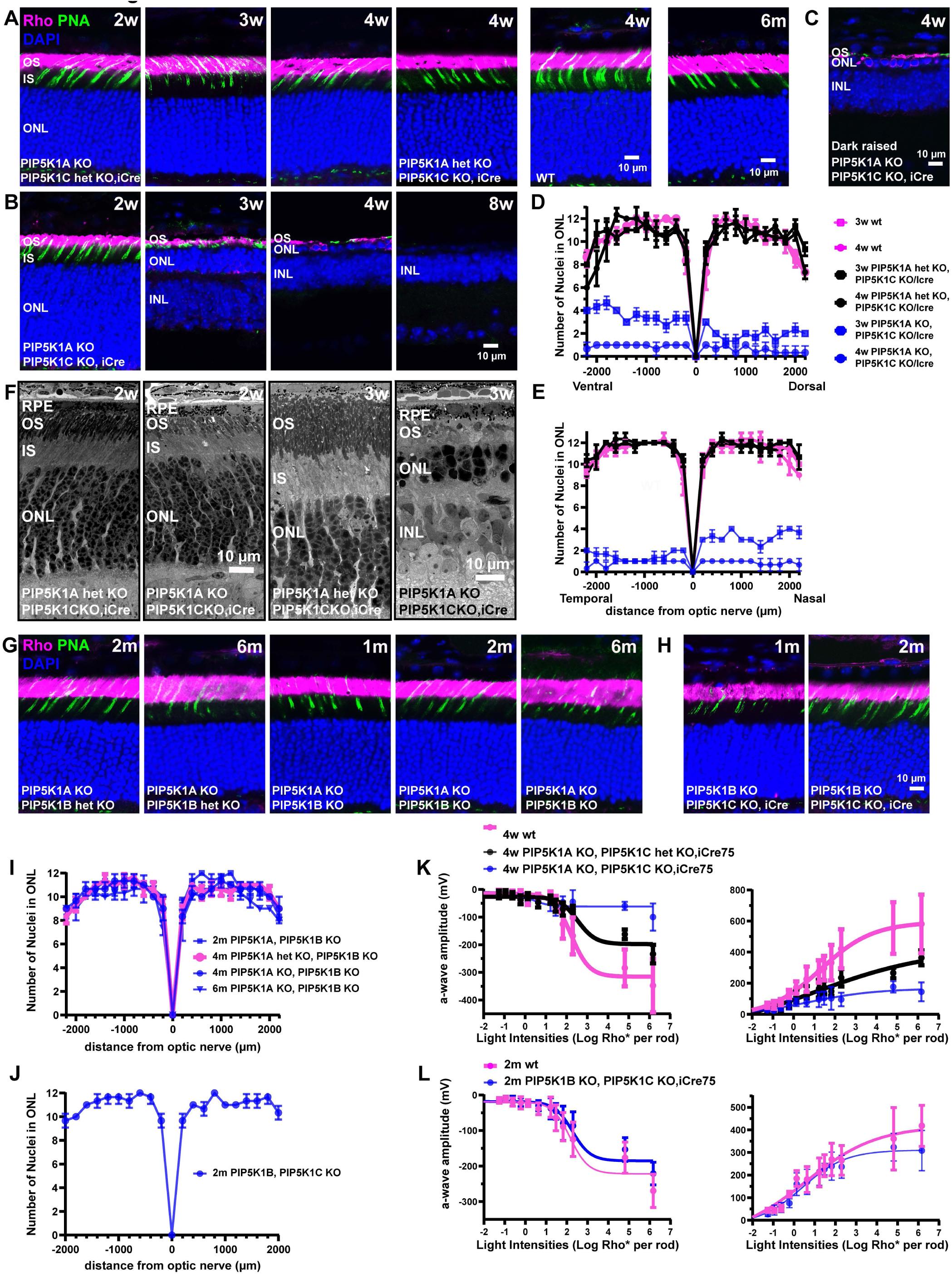
Greatly accelerated retinal degeneration in PIP5K1α + PIP5K1γ double knockout, but not in PIP5K1α + PIP5K1β nor PIP5K1β + PIP5K1γ double knockout. **(A)** Retina sections from PIP5K1α heterozygous knockout **+** PIP5K1 γ knockout (PIP5K1α het KO–PIP5K1γ KO) and PIP5K1α knockout **+** PIP5K1 γ heterozygous knockout (PIP5K1A KO–PIP5K1C het KO) mice, driven by iCre75, were stained with rhodopsin antibody (Rho), peanut agglutinin (PNA, cone sheath marker), and DAPI. No retinal degeneration was observed up to 4 weeks. **(B)** Double knockout of PIP5K1α and PIP5K1γ significantly accelerated retinal degeneration. Photoreceptor cell loss reached ∼50% by 2 weeks and 60–80% by 3 weeks, with no detectable rods by 8 weeks. **(C)** Dark-rearing of PIP5K1α and PIP5K1γ double knockout mice did not prevent retinal degeneration. **(D, E)** “Spidergram” plots show outer nuclear layer (ONL) thickness versus distance from the optic nerve at 3 and 4 weeks. Dorsal and temporal regions of the retina showed somewhat faster degeneration, as compared to more central regions, in 3-week-old PIP5K1α **+** PIP5K1γ double knockout mice. Data are presented as mean ± SEM, *n* = 3 retinas per age group; see also Supplementary Fig. S4. **(F)** Electron micrographs revealed retinal degeneration and disorganized rod outer segments (ROS) in double PIP5K1α and PIP5K1γ knockout mice, without abnormal membrane aggregates were observed. **(G, I)** Double knockout of PIP5K1α and PIP5K1β showed no significant photoreceptor degeneration up to 6 months. **(H, J**; scale bar in **H** applies to **G)** Double knockout of PIP5K1β and PIP5K1γ showed no significant degeneration by 2 months. **(K)** ERG showed ∼50% reduction in a-wave and b-wave amplitudes in PIP5K1α knockout **+** PIP5K1γ heterozygous knockout (PIP5K1A KO–PIP5K1C het KO) mice, despite no visible retinal degeneration (see **A**). Visual responses in PIP5K1α **+** PIP5K1γ double knockouts were nearly absent at 4 weeks. **(L)** No significant changes in ERG responses were observed in PIP5K1β **+** PIP5K1γ double knockout mice at 2 months.

**Figure 8.**
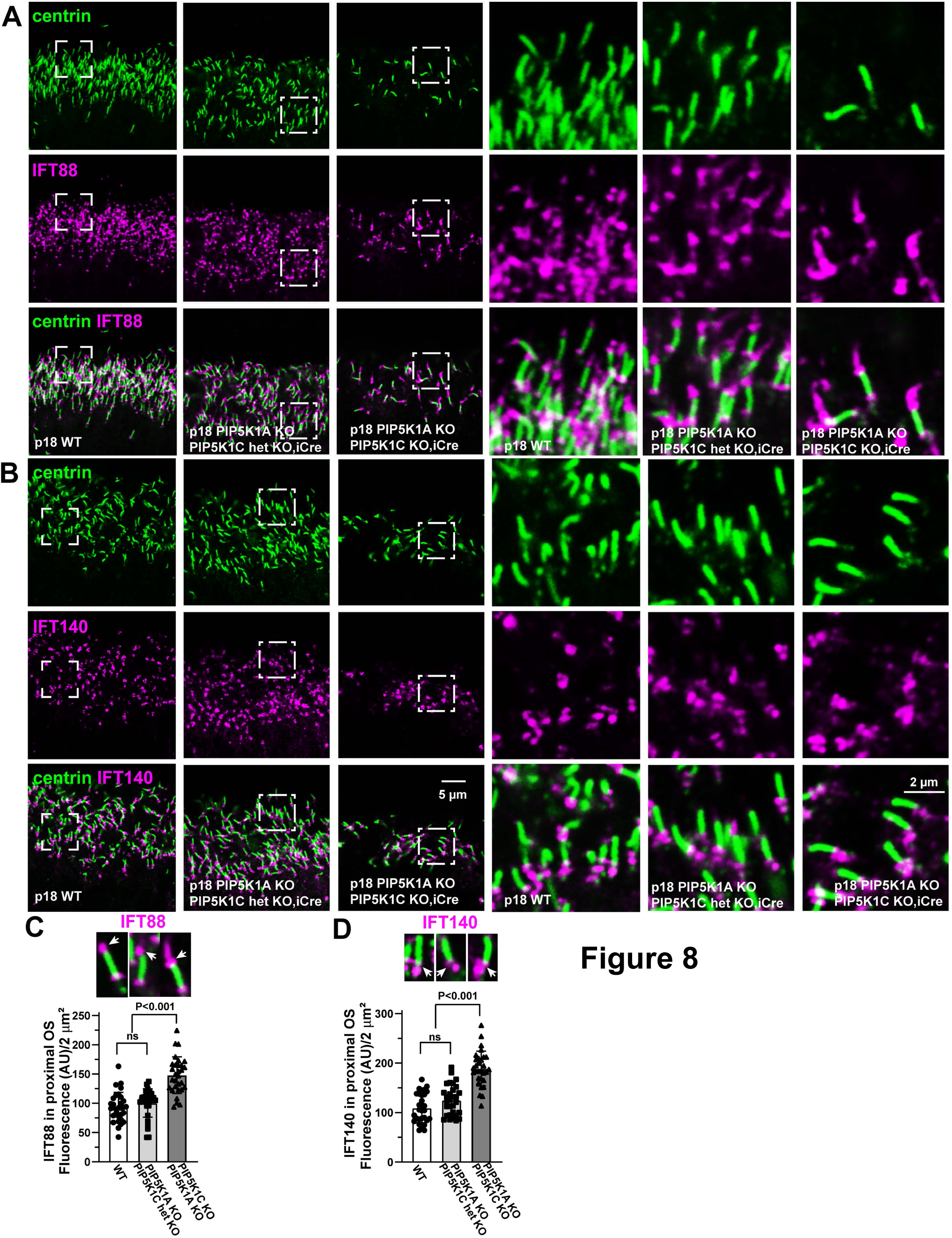
Altered distribution of intraflagellar transport proteins in PIP5K1α and PIP5K1γ double knockout mice. **(A–D)** Immunostaining revealed mis-localization of intraflagellar transport proteins. IFT88 staining extends further into the proximal rod outer segments (ROS) that in WT, and IFT140 accumulated to higher levels near the basal body in PIP5K1α and PIP5K1γ double knockout mice. These findings suggest possible defects in intraflagellar transport. Error bars represent mean ± SEM, *n* = 30 cilia from 3 mice.

**Figure 9.**
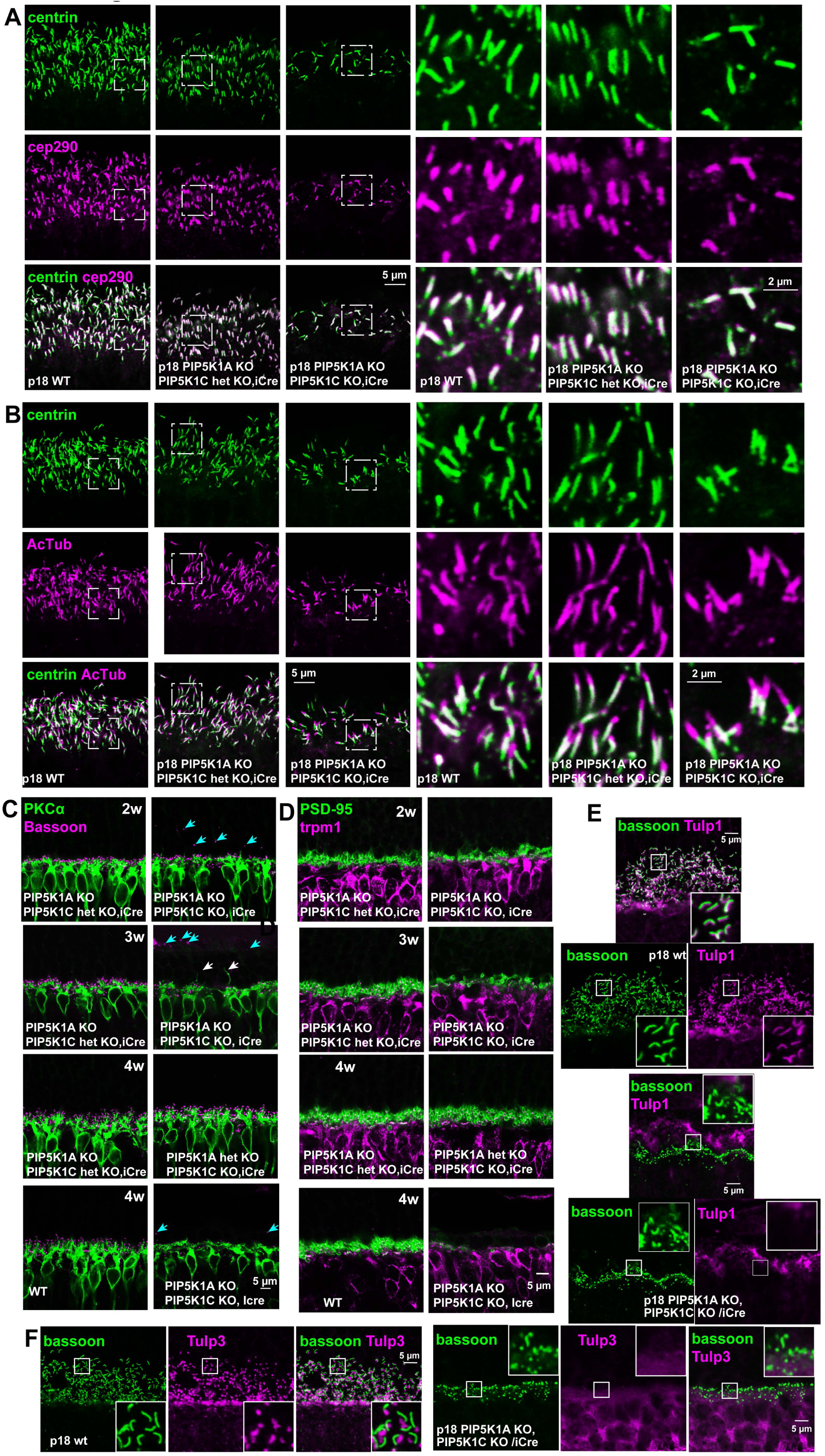
Altered distribution of synaptic proteins in PIP5K1α and PIP5K1γ double knockout mice. **(A, B)** Immunostaining of ciliary proteins centrin, CEP290 and acetylated tubulin, showed distributions in the double knockout mice similar to those in wild-type mice. **(C, D)** Rod bipolar cell dendrites extended into the outer nuclear layer (ONL) at 3 weeks in double knockout mice. Immunoreactivity of photoreceptor synaptic markers Bassoon and PSD-95 was markedly reduced in 4-week-old double knockout mice, consistent with extensive photoreceptor degeneration. **(E, F)** PI(4,5)P_2_-binding proteins Tulp1 and Tulp3, which localize to the wild-type photoreceptor ribbon synapse (marked by Bassoon), lost immunoreactivity in double knockout mice, consistent with depletion of PI(4,5)P_2_ at the photoreceptor synapse. Quantification of alterations in signals for Bassoon, PSD-95, Tulp-1 and Tulp-3, is shown in Supplementary Figure S7.

The most striking results from combined knockouts were observed in mice with both germ-line inactivation of *Pip5k1a* and rod-specific knockout of *Pip5k1c*. Combining germ-line inactivation of *Pip5k1a* with heterozygous rod-specific knockout of *Pip5k1c* did not lead to retinal degeneration or exacerbation of the functional defects seen in the *Pip5k1a* KO (Fig. 7A, K). However, there were abnormalities observed by immunofluorescence, electron microscopy and electroretinography in the case of homozygous rod-specific KO of *Pip5k1c* with homozygous *Pip5k1a* KO (Fig. 7B, C-F, K). This genotype led to greatly accelerated retinal degeneration, relative to either pan-retinal or rod-specific knockout of *Pip5k1c* alone, in the background of WT *Pip5k1a*. At 3 weeks PN there was 66%-90% loss of nuclei in the outer nuclear layer in different regions of the retina, with more temporal and more dorsal regions displaying greater loss (Fig. 7B, D, E, Supplementary Fig. S3). At 4 weeks PN, more than 90% of photoreceptors were lost across the entire retina. Dark rearing (Fig. 7C) did not reduce the rate of degeneration.

Electron microscopy (Fig. 7F, Supplementary Fig. S4) revealed disorganized outer segments, but none of the membrane aggregates or vesicle accumulations seen in some other retinal degeneration genotypes were observed. Basal bodies and connecting cilia appeared normal, despite OS degeneration. At 3 weeks PN, no obvious defects in the outer plexiform layer (OPL) were seen by electron microscopy (Supplementary Fig. S4). Not surprisingly, loss of photoreceptors led to functional defects as observed by ERG (Fig. 7K), with substantial reductions in both a-wave and b-wave amplitudes at 4 weeks PN.

To explore the effect of the knockouts on protein localization in rods, we used immunofluorescence labelling of proteins specific for the connecting cilium (Figs. 8, 9A-B) or for photoreceptor synaptic regions (Fig. 9C-F). Spatial distribution of the connecting cilium proteins, acetylated tubulin, centrin and CEP290 appeared normal (Figs. 8A, B, 9A, B).

Two proteins of the intraflagellar transport machinery, IFT140 and IFT88, displayed abnormal accumulation in the connecting cilium (Fig. 8A-D). In immunofluorescence procedures carried out and imaged in parallel using identical conditions, at 18 days PN, IFT40 fluorescence in the connecting cilia was 75% (± 33%) higher in the double knockout (*Pip5k1a* KO plus rod-specific *Pip5k1c* KO) animals than in WT, and IFT88 fluorescence was 58% (± 34%) higher. These results suggest a defect in retrograde transport of the IFT complex components, and possibly defects in ciliary trafficking more generally, as the IFT complexes play a critically important role in ciliary trafficking. Electron microscopy at 2 and 3 weeks PN (Supplementary Fig. S4A, B) revealed no obvious alterations in CC structure at either age, but severe disruption of OS structure at 3 weeks in the *Pip5k1a* KO plus rod-specific *Pip5k1c* KO retinas.

### Synaptic layer defects in α/γ double KO

In the outer plexiform layer (OPL, Fig 9C-D), we used PKCα and TRPM1 as markers of ON bipolar cells in sections prepared from double knockouts. PKCα labels the cell bodies and dendrites, but not the dendritic tips of rod bipolar cells, whereas TRPM1 labels the dendritic tips, dendritic shafts and intracellular membranes throughout both rod and cone ON-bipolar cells (Agosto et al., 2018b; Agosto et al., 2018a). For photoreceptor terminals, we used synaptic ribbon marker, Bassoon, and pre-synaptic plasma membrane marker PSD95 (Fig 9C-F). In addition, we probed with antibodies specific for the phosphoinositide binding proteins TULP1 and TULP3 (Fig 9E-F). As early as 2 weeks PN, the double knockout retinas (*Pip5k1a* KO plus rod-specific *Pip5k1c* KO) had signs of photoreceptor synaptic termini retraction (Fig. 9C, cyan arrows) and by 3 weeks PN displayed bipolar cell sprouting (Fig. 9C, white arrows), as frequently seen in retinas experiencing photoreceptor cell death. By 2 weeks PN, PSD95 staining (Fig. 9D) was already more diffuse in the double knockouts than in WT, with much less of the characteristic U-shaped staining of this pre-synaptic plasma membrane marker. At 3 weeks PN, PSD95 signal was weaker as well as displaying aberrant spatial distribution, and by 4 weeks PN, the staining had largely disappeared. A similar pattern was observed for Bassoon staining, with altered morphology by PN day 18 and declining signal at later stages (Fig. 9C,E-F), with almost complete loss by 4 weeks PN. Quantification of alterations in synaptic markers is shown in Supplementary Fig. S7.

The phosphoinositide-binding protein, TULP1 is expressed in photoreceptor cells and is readily detected by immunostaining in the OPL ((Ikeda et al., 1999; Grossman et al., 2009) and Fig. 9E). Single cell RNA sequencing data (Clark et al., 2019) confirm high levels in both rods and cones. It displays a distinct pattern of staining immediately adjacent to the ribbon staining for Bassoon and parallel to it across the arc of the ribbon. TULP3 RNA is expressed in postsynaptic cells in the OPL (Ikeda et al., 1999). TULP3 immunostaining is similar to but distinct from TULP1 staining (Fig. 9F). It has a more punctate pattern, close to, but not immediately adjacent to the center of the ribbon arc (Fig. 9F), and co-localized with TRPM1 staining in ON-bipolar cell dendritic tips. By PN day 18, the characteristic WT staining patterns adjacent to the Bassoon crescents were lost for both TULP1 and TULP3, indicating a critical need for PI(4,5)P_2_ supplied by PIP5KIα and/or presynaptic PIP5KIγ for their proper localization. Quantification of alterations in TULP1 and TULP3 staining is shown in Supplementary Fig. S7. CC staining for TULP1 was detected in only a small fraction of cells even in WT retina, as reported previously (Sharif et al., 2021), and could not be reliably distinguished from background puncta.

Despite the alterations in distributions of some OPL marker proteins, electron microscopy of the α/γ double-KO at 2 or 3 weeks postnatal (Supplementary Fig. S4C, D) did not show obvious alterations in synaptic structures, such as the ribbons, nor was accumulation of vesicles observed. There were fewer ribbons apparent in micrographs at 3 weeks PN (Supplementary Fig. S2D), as expected from the greatly reduced numbers of photoreceptors at this age in the double knockouts, but our TEM data do not provide evidence for direct effects of the double knockout on synapse ultrastructure.

PI(4,5)P_2_ has been reported to play an important role in vesicle exocytosis (Martin, 2012), so we examined immunostaining patterns for two proteins important in this process in the retina (Dey et al., 2024), syntaxin3 (Stx3, whose b splice variant is retina-specific) and VAMP2, an essential component of synaptic vesicles (Supplementary Fig. S5A). At 18 d PN in the PIP5KIα plus rod-specific PIP5KIγ KO, signal intensities were not significantly lower for Stx3, and only slightly lower (*p* = 0.038) for VAMP2 (Supplementary Fig. 5C. D). The distribution of Stx3 signal was significantly narrower in the double knockouts (*p* < 0.003, Fig. 5C, D), but not that of VAMP2. At 7 weeks PN in the PIP5KIα plus Six3-Cre-driven PIP5KIγ double KO retinas, there was extensive accumulation of large VAMP2-containing clumps and total VAMP2 signal was reduced by nearly one-half (quantification shown in Supplementary Fig. S7).

In contrast, there were no obvious changes by immunostaining in levels of transferrin, which is imported into photoreceptors by endocytosis (Supplementary Fig. S5B), in endosome markers EEA1, Rab5 and Rab7 (Supplementary figure S6) nor in localization to the distal region of rods of the α subunit of the cyclic nucleotide-gated channel, CNGA1, which is normally localized to rod OS by vesicular transport (Supplementary Fig. S5B). Transport of the rod G protein, transducin (Gtα) and its effectors, the cGMP-gated channel, to OS did not appear to be substantially diminished in the α/γ double knockouts (Supplementary Figures S5, S6).

Although we did not explore cell death pathways extensively, we did check for evidence of enhanced autophagy in mice with germ-line inactivation of *Pip5k1a* combined with heterozygous rod-specific knockout of *Pip5k1cγ* by immunostaining for markers of autophagy, p62 and LAMP1 (Supplementary Fig. S6C). We did not see the accumulation of these proteins that we observed in a positive control for perturbations of autophagy, rod-specific Vps3 KO (Supplementary Fig. S6C).

We also examined distributions of some of the CC and OPL proteins in other double knockouts (Fig. 10). Even at 6 months PN, double knockouts of *Pip5k1a* plus *Pip5k1b* displayed no evidence of abnormal ciliary or synaptic protein labeling. Signal for CC proteins, centrin, CEP290, IFT140, and IFT88 were not noticeably different from controls (Fig. 10 A-C). Likewise, patterns for OPL markers Bassoon and PKCα (Fig. 10D) appeared normal.

**Figure 10.**
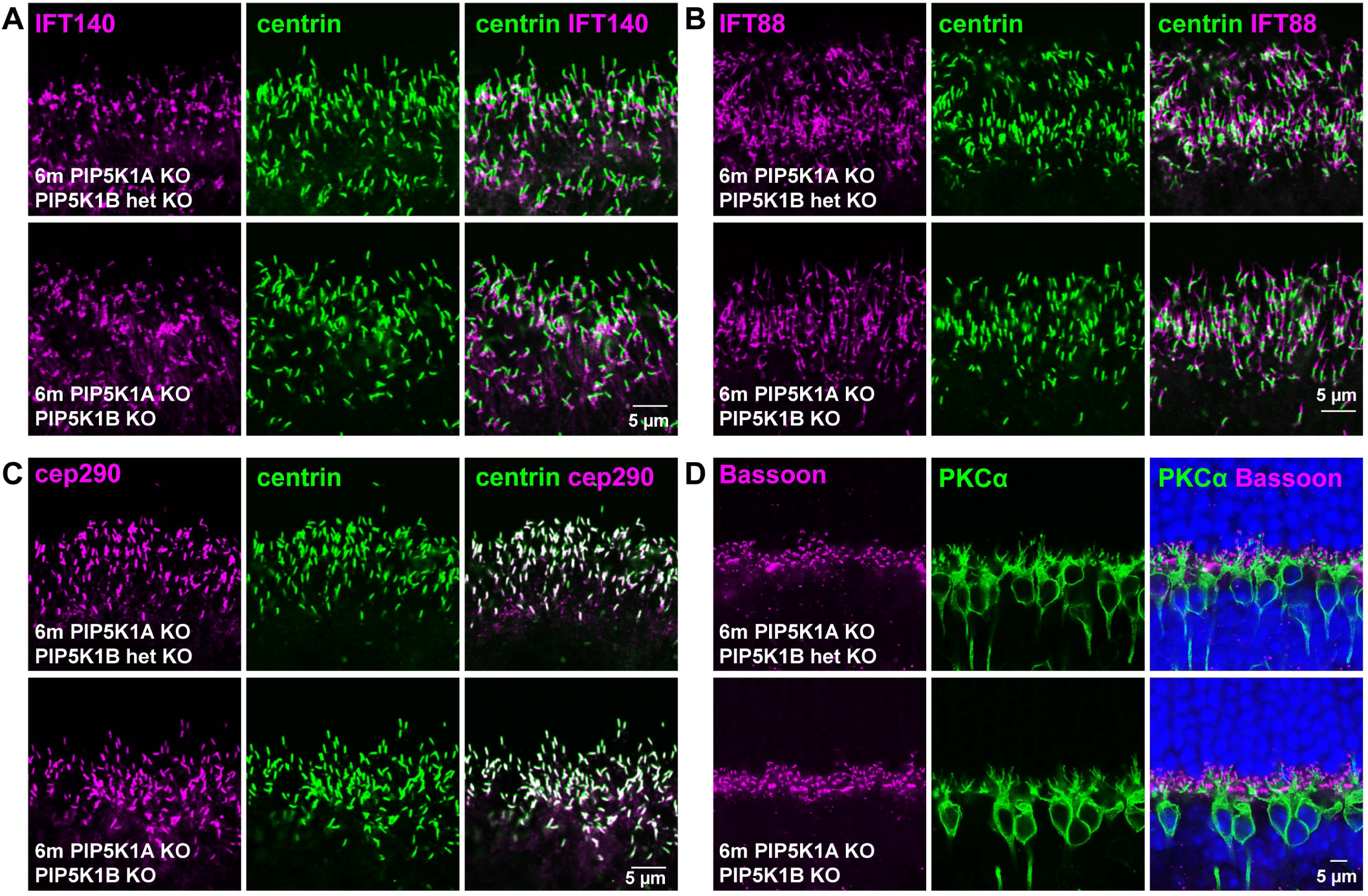
No obvious morphology changes in connecting cilia and ribbon synapses of PIP5K1A and PIP5K1B double knockout photoreceptor cells. **(A, B)** immunostaining centrin with intraflagellar transport proteins IFT88 and IFT140 showed similar distribution in PIP5K1A KO **+** PIP5K1B het KO and PIP5K1A **+** PIP5K1B double KO. **(C, D)** Ciliary proteins centrin, CEP290, and ribbon marker Bassoon showed without noticeable difference between PIP5K1A KO-PIP5K1B het KO and PIP5K1A-PIP5K1B double KO.

### Effects of Triple knockouts

Although the combined knockouts of all three kinases had greatly reduced fertility, we managed to breed some animals whose genotype included *Pip5k1a^-/-^, Pip5k1b^-/-^,*and *Pip5k1c^fl/f/iCre75^*. At 2 weeks after birth, photoreceptor degeneration was already obvious, with a 20% reduction in the number of nuclei in the ONL and sprouting of bipolar cell dendrites (Supplementary Fig. S8), indicating that additional loss of PIP5K1β function further exacerbates the phenotype of the *Pip5k1a Pip5k1c* double KO in rods. Thus, despite the negligible effects of knocking out *Pip5k1b* by itself, or together with knockouts of either *Pip5k1a*, *Pip5k1c*, PIP5KIβ is able to fulfill some of the functions of the α and γ isoforms when both are missing.

## Discussion

The results of this work reveal the critical importance of PI(4,5)P_2_ for photoreceptor function, health, and survival in the mammalian retina. This conclusion, while novel, is perhaps not too surprising given the ubiquitous importance of PI(4,5)P_2_ in metazoan biology.

The initial finding that light-adapting conditions lead to a dramatic increase in PI(4,5)P_2_ levels in isolated rod cell fragments enriched in rod outer segments (ROS, Fig. 1A, Supplementary Fig. S1) suggested the hypotheses that PI(4,5)P_2_ may be present in rod outer segments, that its levels there may be modulated by phototransduction, and that it may, in turn, modulate phototransduction. The presence of PI(4,5)P_2_ in ROS was subsequently confirmed (Fig. S1D) as was its dependence on phototransduction (Fig. 1B).

While there are no results demonstrating modulation of phototransduction by PI(4,5)P_2_ in response to light, the decreases in physiological responses observed in the case of the PIP5KIα KO suggest that one or more steps in the phototransduction pathway is modulated by the local pool of PI(4,5)P_2_ generated by this isoform. It is unlikely that the steps leading from light activation of rhodopsin to depletion of cGMP, which transpire predominantly on disc membranes, are directly modulated by PI(4,5)P_2_, which is localized to the plasma membrane. However, cyclic nucleotide-gated (CNG) channels, which are localized to the same membranes in outer segments as PI(4,5)P_2_, have been previously reported to be modulated by PI(4,5)P_2_ (Womack et al., 2000), and there is evidence for phosphoinositide modulation of multiple CNG channels as well as cyclic nucleotide-regulated HCN channels, another type of channel that shapes electrical responses of vertebrate photoreceptors and bipolar cells ((Hilgemann et al., 2001; Demontis et al., 2002; Müller et al., 2003; Brady et al., 2006; Bright et al., 2007; Pian et al., 2007; Barrow and Wu, 2009; Flynn and Zagotta, 2011; Ying et al., 2011; Ramakrishnan et al., 2012; Dai et al., 2013; Dai and Varnum, 2013; Dai et al., 2014; He et al., 2014; Sothilingam et al., 2016; Ingram et al., 2020; Claveras Cabezudo et al., 2022; Pang et al., 2024; Thon et al., 2024; Hartveit et al., 2025). Further electrophysiological studies will be needed to elucidate the roles of PIP5KI isozymes and PI(4,5)P_2_ in retinal synaptic signaling.

Given the death of rods caused by deficiencies in the γ and α isoforms of PIP5KI, it seems likely that the light-driven increases in PI(4,5)P_2_ are also involved in homeostatic mechanisms that promote cell survival during prolonged exposure to bright light of these exquisitely light-sensitive cells. The accumulation of intracellular transport proteins in the connecting cilium in the double *Pip5k1a* KO plus rod-specific *Pip5k1c* KO retinas suggest that regulation of ciliary transport may be one such homeostatic mechanism.

Both the relatively slow progression of cell death in the case of rod-specific PIP5KIγ KO and the early onset nature of the double α/γ KOs present challenges for determining mechanisms of cell death. There are few obvious morphological changes preceding cell death in the case of the single PIP5KIγ KO, and the overlap between the completion of photoreceptor development and cell death onset in the double knockout make it difficult to resolve processes that cause defective OS development, such as IFT protein accumulation, from pathological events that occur after maturation. One clue may be the axon retraction observed in both rod-specific PIP5KIγ KOs and the double knockouts. Such retraction is a typical response to injuries, such as retinal detachment (Fisher et al., 2005; Linberg et al., 2009), and has also been observed in response to genetic deficiencies in post-synaptic cells, such as mGluR6 deficiency (Bayley and Morgans, 2007) or horizontal cell ablation (Sonntag et al., 2012). An example of rod axon retraction has been reported in a model of photoreceptor degeneration, canine deficiency in the X-linked *RPGR* gene, whose deficiency in mice we recently reported to be linked to changes in actin dynamics (Megaw et al., 2024), and in dogs was observed to be linked both to actin dynamics and decreased expression of the axon-guidance receptor, ROBO1 (Appelbaum et al., 2020). The persistence of substantial numbers of retracted axons in every field we imaged from the mouse KOs suggest that this major change in axonal structure precedes cell death. However, more experiments will be needed to determine whether these changes contribute to cell death or are caused by more fundamental changes that accompany cell death.

Some conclusions from our results are quite surprising. One of these is the apparent *lack* of importance of the PIP5KI family in retinal development. While this conclusion is robust with respect to the α and β isoforms, as those were tested with germ-line knockouts, with respect to the γ isoform it is possible that some essential developmental functions may be masked by the incomplete nature of the Six3-Cre-driven *Pip5k1c* knockout and the late-stage (PN day 10-18) nature of the iCre75-driven rod-specific *Pip5k1c* knockout. It may be confidently concluded, however, that PI(4,5)P_2_ and the PIP5KI enzymes do not play a key role in the formation of rod outer segments and their associated ciliary structures, but are essential for maintaining the outer segment structure.

Also somewhat surprising is the remarkable degree of redundancy that seems to be built into the PI(4,5)P_2_ synthesis pathway. Clearly each of the three isoforms can fulfill parts of the essential functions of the others when they are missing, as revealed by the exacerbation of the phenotypes of single knockouts by pairing with additional knockouts (*i.e.*, adding *Pip5k1a* or *Pip5k1c* knockout to any single knockout) and the exacerbation of the *Pip5k1a*/*Pip5k1c* knockout by adding *Pip5k1b* knockout. Whereas rod-specific *Pip5k1c* knockout is the only single-isoform knockout sufficient to cause retinal degeneration on its own, *Pip5k1a* knockout is sufficient to reduce phototransduction efficiency without inducing morphological changes. Moreover, both the *Pip5k1a* and *Pip5k1b* knockouts worsen the *Pip5k1c-*dependent rod degeneration phenotype when combined with others. These observations are reminiscent of previous observations that *Pip5k1a* alone, but not *Pip5k1b*, is sufficient to support prenatal development in the absence of functional *Pip5k1c,* whereas *Pip5k1b* KO substantially worsens the postnatal phenotype of the *Pip5k1c* KO (Volpicelli-Daley et al., 2010).

The mechanisms for apparent redundancy remain to be determined. We have some evidence for increases in PIP5KIα protein levels in the *Pip5k1c* KO (Fig. 3F), but additional experiments will be needed to test for compensatory upregulation of activities of individual isoforms in knockout retinas. Another possibility to consider would be altered localization of one isoform in cells missing another isoform.

Although the highest concentrations of PI(4,5)P_2_ in the outer retina are in the synaptic region, nothing in the results presented here supports an essential role for this lipid or the kinases that produce it in synaptic function. However, the scotopic electroretinogram is primarily sensitive to rod-to-bipolar cell synapses, and is a pooled measurement that could miss subtle changes at the levels of individual cells, kinetics of vesicle transport and recycling, *etc.* Certainly, there are signs of synaptic irregularities accompanying PIP5KI KOs in the outer retina, such as alterations in the subcellular distributions of Stx3 and Vamp2 in *Pip5k1c* KOs, consistent with a previous report that the γ isoform is the major PI(4,5)P_2_-enzyme at synapses in the brain (Wenk et al., 2001). Distribution of the PI(4,5)P_2_-binding proteins, TULP1 and TULP3 in the vicinity of synapses is also disrupted by PIP5K1 deficiencies, but their roles in synaptic function are not well understood. TULP1 KO has been shown to lead to synaptic defects in the outer retina (Grossman et al., 2009). Quantitative analysis of the perturbations observed in these synaptic markers is provided in Supplementary Fig. S7.

The PI(4,5)P_2_-binding proteins, TULP1 and TULP3 have also been implicated in cilium function, with TULP3 reported to link IFT complex trafficking to PI(4,5)P_2_ (Mukhopadhyay et al., 2010), and the abnormal accumulation we observe of proteins of both the IFTA (IFT140) and IFTB (IFT88) subcomplexes is consistent with an important role for PI(4,5)P_2_ in IFT trafficking.

The enrichment of PI(4,5)P_2_, of both PIP5K1α and PIP5k1γ, and of PI(4,5)P_2_ - binding proteins TULP1 and TULP3 in the synaptic region of the outer plexiform layer, and the perturbation of distributions of synaptic marker proteins in the knockout mice support a role for PI(4,5)P_2_ and PIP5K1 activity in synaptic function. We have no direct evidence supporting such a function, but it would be consistent with previous studies of their roles at other synapses in the nervous system.

Genetic evidence (Verstreken et al., 2003; Van Epps et al., 2004; George et al., 2014) supports a role in synaptic signaling in *Drosophila* and zebrafish retina for the dual-specificity phosphoinositide phosphatase synaptojanin1, which can catalyze dephosphorylation of PI(4,5)P_2_, and plays a major role in synaptic vesicle recycling (Cremona et al., 1999; Chang-Ileto et al., 2011; De Camilli, 2026). In mice PI(4,5)P_2_ and PIP5k1γ play a major role in synaptic membrane recycling in the brain (Wenk et al., 2001; Di Paolo et al., 2004). This role is likely linked to the importance of PI(4,5)P_2_ in regulating actin cytoskeletal reorganization (El Sayegh et al., 2007; Mao and Yin, 2007; Fairn et al., 2009; Zhang et al., 2012; Hansen et al., 2022; Calabrese and Halpain, 2024; Albanesi et al., 2026). PI(4,5)P_2_ has been reported to play a role in synaptic vesicle docking in hippocampal neurons (Chen et al., 2021) and in regulating the readily-releasable vesicle pool in chromaffin cells (Milosevic et al., 2005). In post-synaptic cells PI(4,5)P_2_ has been reported to play a role in membrane bending and dendritic spine formation (Hotulainen and Saarikangas, 2016).

As in many slowly progressing retinal degenerations, the mechanisms underlying photoreceptor cell death in all genotypes lacking functional PIP5K1γ remains unclear. Likewise, the exact sequence of events leading to disrupted OS morphology remains to be determined, although the IFT protein accumulations suggest defects in ciliary trafficking are likely involved. Further investigations of these defects, as well as those underlying reduced ERG amplitudes, will be important subjects for future study.

## Materials and Methods

### MICE

All animal procedures were approved by the Institutional Animal Care and Use Committee at Baylor College of Medicine and conducted in accordance with NIH guidelines and the ARVO Statement for the Use of Animals in Ophthalmic and Vision Research. Wild-type C57BL/6J (Strain #000664) and B6.129S4-Gt(ROSA)26Sortm1(FLP1)Dym/RainJ (Strain #009086) mice were purchased from Jackson Laboratory (Bar Harbor, ME), and CD-1 IGS mice from Charles River (Wilmington, MA). The Gnat1 (rod transducin alpha) knockout mice was kindly given by Dr. Janice Lem, Tufts University (Calvert et al., 2000). The knockout-first allele (tm1a) of *Pipk1c* mice (C57BL/6N-Pip5k1ctm1a(KOMP)Wtsi/JMmucd) was purchased from the Mutant Mouse Regional Resource Center (MMRRC), University of California, Davis. These mice were crossed with FLP-expressing mice to excise the LacZ-neomycin cassette, generating a *Pipk1c* floxed line. *Pipk1c* conditional knockout mice were produced by breeding *Pipk1c* floxed mice with mice bearing a Cre transgene, either iCre75 (gift from Dr. Ching-Kang Jason Chen, Baylor College of Medicine; Genesis 2005, 41(2):73–80) or Six3-Cre (Tg(Six3-cre)69Frty/GcoJ (Furuta et al., 2000) gift from Dr. Melanie Samuel, Baylor College of Medicine) mice to achieve photoreceptor- or retina-specific *Pipk1c* deletion. The sperm from PIP5K1A knockout mice (B6;129S5-Pip5k1aGt(OST43713)Lex/Mmucd, stock #011756-UCD) and *Pipk1b* knockout mice (B6.129P2-Pip5k1b<tm1Tssk>, RBRC10384) were obtained from MMRRC, and RIKEN BioResource Research Center (Tsukuba, Japan), respectively. In vitro fertilization (IVF) of C57BL/6J oocytes with these sperm was performed at the Genetically Engineered Rodent Models Core, Baylor College of Medicine, to generate *Pipk1a* and *Pipk1b* knockout lines. Tail DNA was used for *Pipk1* allele genotyping following the instructions from the companies. The absence of *Pde6brd1* was confirmed by PCR; absence of Crb1rd8 was confirmed by PCR and sequencing of the relevant Crb1 region. For *RD10* mutations, PCR primers 5’-GTGTGAGAGCCCTTCCCAAACAGG-3’ and 5’- GGTGTCCCAAACCCATCCCTTTG-3’ were used to amplify exon13 of *Pde6b*, primers 5’- CAGGACACTGACAGAGGGTAG-3’ and 5’- GGAGTAGGTGACAGACTGGCC-3’ for amplifying exon 16 of *Pde6b*. The PCR products were sequenced to confirm absence of RD10 mutations (Chang et al., 2007). Litter sizes were near normal in single isoform of PIP5 kinase knockouts (*Pipk1a* KO, *Pipk1b* KO, *Pipk1c* conditional KO), as well as in double knockouts (*Pipk1a* KO– *Pipk1c* conditional KO and *Pipk1b* KO– *Pipk1c* conditional KO). However, significantly reduced litter sizes (1-3 pups) were observed in mating pairs of *Pipk1a* homozygous KO– *Pipk1b* heterozygous KO and *Pipk1a* heterozygous KO– *Pipk1b* homozygous KO. Mice used for tissue collection were euthanized by CO₂ inhalation prior to dissection, following protocols from the American Association for Laboratory Animal Science.

### ANTIBODIES

The antibodies and reagents used in immunoblotting and immunostaining are: Rabbit anti-PIP5KIA (10092-348, Proteintech); Rabbit anti-IFT140 (17460-1-AP, Proteintech); Rabbit anti-IFT88 (13967-1-AP, Proteintech); Rabbit anti-TULP1 (18971-1-AP, Proteintech); Rabbit anti-TULP3 (13637-1-AP, Proteintech); Rabbit anti-PIP5KIB (gifted by Dr. Yuji Funakoshi, University of Tsukuba, Japan); Rabbit anti-PIP5KIC (3296S, Cell Signaling); Rabbit anti-PKCα (2056S, Cell Signaling); Rabbit anti-GAPDH (D16H11) XP® (5174S, Cell Signaling); Rabbit anti-LC3A/B (D3U4C) (12741S, Cell Signaling); Rabbit anti-Rab5 (C8B1) mAb (3547S, Cell Signaling); Mouse-anti-EEA1 (E9Q6G) (48453S, Cell Signaling); Rabbit anti-Cre (69050-3, EMD-Millipore); Mouse anti-Lamp1 mAb (LY1C6) (428017, Millipore-Sigma); Mouse anti-Centrin, clone 20H5 (04-1624, Millipore-Sigma); Rabbit anti-CNGA1(ab136123, Abcam); Mouse anti-CNGA1 (ab253296, Abcam); Rabbit anti-Rab7 [EPR7589] (ab137029, Abcam); Rabbit anti-Lamp1(ab30687, Abcam); Rabbit anti-ribeye (192 103, YSYS); Peanut Agglutinin (PNA)-Rhodamine (RL-1072-5, Vector Laboratories); Mouse anti- PKCα (H-7) (sc-8393, Santa Cruz Biotechnology); Rabbit anti-Na+/K+-ATPase α (H-300) (sc-28800, Santa Cruz Biotechnology); Rabbit anti-α Tubulin Antibody (H-300) (sc-5546, Santa Cruz Biotechnology); Mouse anti-acetylated alpha Tubulin (6-11B-1) (sc-23950, Santa Cruz Biotechnology); Mouse anti-Bassoon Antibody (SAP7F407) (sc-58509, Santa Cruz Biotechnology); Goat anti-Calbindin D28K (C-20) (SC-7691, Santa Cruz Biotechnology); mouse anti-PSD-95 (K28/43) (75-028, NeuroMab); Rabbit anti-Rhodopsin (GTX129910, GeneTex); Guinea pig anti-p62, C-Terminal (03-GP62-C, American Research Products); Rabbit anti-Cep290 (A301-659A, Bethyl Laboratories); Mouse anti-Rhodopsin (1D4) (prepared in-house from hybridoma culture medium); Mouse anti-Trpm1 (274G7) (prepared in-house from hybridoma culture medium); DAPI (D9542, Sigma). ProLong™ Diamond Antifade Mounting Medium (P3961, Invitrogen); VECTASHIELD Antifade Mounting Medium with DAPI (H-1200, Vector Laboratories).

### ELECTRORETINOGRAPHY (ERG)

The mice were dark-adapted overnight prior to electroretinography (ERG). Under dim red light, they were anesthetized with a solution of ketamine (95 mg/ml) and xylazine (5 mg/ml). A single drop each of 1% tropicamide and 2.5% phenylephrine was applied to dilate the pupils. The mice were then placed on a heating pad maintained at 39 °C inside a Ganzfeld dome coated with reflective white paint. A small amount of 2.5% methylcellulose gel was applied to each eye, and a platinum electrode was positioned in contact with the center of the cornea. Reference and ground platinum electrodes were placed on the forehead and tail, respectively.

After placement in the dome, the mice remained in complete darkness for several minutes. Signals were amplified using a Grass Instruments amplifier (Quincy, MA) with a bandpass filter of 0.1 to 1,000 Hz. Data were acquired using a National Instruments Lab-PC data acquisition board at a sampling rate of 10,000 Hz (National Instruments, Austin, TX). The traces were averaged and analyzed using custom software written in MATLAB (MathWorks, Natick, MA).

### SUBRETINAL INJECTION AND ELECTROPORATION

These were carried out as initially described (Matsuda and Cepko, 2004, 2007) and later modified (He et al., 2016). The pE-7T-E construct (Lee et al., 2010), was generously gifted by Dr. Joseph C. Corbo (Washington University in St. Louis). It contains a single mutation in the second Crx transcription factor binding site of the mouse opsin promoter. This mutation reduces mouse opsin promoter activity by over 90%, helping to mitigate the issues related to target protein overexpression in transfected photoreceptor cells. GFP cDNA, GFP-2xFAPP1 cDNA that contained 2 copies of amino acids 1-100 of human four-phosphate-adaptor protein 1 PH domain, and DsRed-PLCδ cDNA containing amino acids 1–175 of human phospholipase C δ1 PH domain (He et al., 2025) were subcloned into the EcoRI and NotI sites of the pE-7T-E vector. Subretinal injections and electroporation were used to deliver the E-7T-E mutant opsin promoter constructs (DsRed-PLCδ or GFP-2xFAPP1 plasmid DNA at 2 mg/ml) into photoreceptor cells of P0 wild-type CD1 mouse pups. The lack of pigment in CD1 mice facilitates visualization during injection. After anesthesia, an incision was made at the future eyelid margin, followed by creation of a pilot hole using a 30-G beveled needle. A 33-G blunt needle was then positioned into the subretinal space. Approximately 450 nl of plasmid DNA in PBS containing 0.1% Fast Green dye was injected using a UMP3 Microsyringe Injector and Micro4 Controller (World Precision Instruments). Electroporation was performed using five square-wave pulses (80 V, 50 ms duration, 950 ms intervals) delivered by an ECM 830 electroporator (BTX Harvard Apparatus) and custom tweezer-type electrodes, with the negative electrode placed on the injected eye. Injected eyes were harvested for imaging at four weeks post-injection.

### TRANSMISSION ELECTRON MICROSCOPY (TEM)

TEM was performed as previously described (He et al., 2019; He et al., 2025). Eye cups were fixed in a solution containing 3% paraformaldehyde and 3% glutaraldehyde in 0.1 M cacodylate buffer (pH 7.4) for 2 days at 4 °C. After fixation, the samples were washed and post-fixed with 1% osmium tetroxide (OsO₄) in the same buffer for 1 hour. Dehydration was carried out through a graded ethanol series (30%, 50%, 70%, 85%, 90%, and 100%), followed by multiple acetone washes. Tissues were gradually infiltrated with acetone:resin (Embed-812, Electron Microscopy Sciences, Hatfield, PA) mixtures in ratios of 2:1, 1:1, and 1:2, then immersed in pure resin for 48 hours. Polymerization of resin blocks occurred at 60 °C for 24–48 hours. Ultrathin sections (80–90 nm) were cut using a Leica Ultracut UCT microtome, mounted on 50 or 75 mesh grids, and stained with 2% uranyl acetate and Reynold’s lead citrate. Imaging was performed using a Zeiss transmission electron microscope equipped with a CCD camera (Advanced Microscopy Techniques Corp., Woburn, MA).

### IMMUNOSTAINING

To establish the orientation of the mouse eye following euthanasia, a burn mark is applied to the dorsal cornea. This technique, adapted from (Sondereker et al., 2018), involved heating an 18-gauge stainless steel syringe needle using a gas burner and briefly touching the tip to the dorsal cornea for less than one second. The resulting mark is positioned directly between the nasal and temporal regions, providing a reliable anatomical reference for subsequent dissection and analysis.

Mouse eyes were excised and rinsed with PBS, then fixed in 4% paraformaldehyde in PBS for 45 minutes at room temperature. For cryosectioning, mouse eyecups were cryoprotected overnight at 4 °C in 26% sucrose in PBS and embedded in optimum cutting temperature (O.C.T.) compound. The mouse eyecups were sectioned using Thermo HM525 NX cryostat. Eyecup sections were washed with PBS, then incubated in blocking solution (10% donkey serum, 5% BSA, 0.4% fish gelatin, and 0.4% triton X-100 in PBS) at room temperature for 1 hour. Samples were then incubated overnight at room temperature with primary antibodies diluted 1:50–1:100 in blocking solution. After PBS washes, secondary antibodies conjugated with 2 µg/ml Alexa dye (Thermo) and 300 nM DAPI (Sigma) were applied in blocking solution for 1 hour at room temperature. Samples were washed again with PBS and mounted using VECTASHIELD (Vector Laboratories) or ProLong™ Diamond antifade mounting medium (Invitrogen). Samples were imaged using a Leica TCS SP5 confocal microscope equipped with a Leica 63x oil immersion objective (HCX PL APO CS 63.0x, NA 1.40 oil) and lasers including diode 405, argon 488, HeNe 543, and HeNe 633. Tiled images of entire retinal sections were acquired using a Zeiss LSM 710 confocal microscope system (Zeiss, Oberkochen, Germany) with a 40x water immersion objective (Zeiss C-Apochromat 40x/1.2W Korr UV-VIS-IR). Each tile measured 1024 × 1024 pixels, and a total of 289 tiles (covering 3613.32 × 3613.32 µm) were stitched together using ZEN 2 (blue edition) software (Zeiss). ImageJ was used for converting the original images to tiff files and for quantification (see *Data Analysis*, below). For figure preparation, contrast and brightness were adjusted in Photoshop (Adobe).

For immunostaining the connecting cilia markers, the fresh unfixed mouse eyes were embedded into O.C.T. compound, and sectioning. The mouse eye sections were collected on poly-lysine coated slides (Electron Microscopy Sciences) and fixed on the slides with 1% paraformaldehyde in PBS for 2 minutes at room temperature. The immunostaining was performed as above after post-fixation.

### IMMUNOBLOTTING

Two mouse retinas were suspended in 160 µl RIPA buffer containing protease inhibitor cocktail and sodium orthovanadate (Santa Cruz Biotechnology, Inc.) and sonicated on ice for 2 minutes. The retinal lysate (10–15 µl) was mixed with 2x Laemmli Sample Buffer (Bio-Rad) and separated by SDS-PAGE. After transferring to nitrocellulose membranes, the blots were blocked with EveryBlot Blocking Buffer (Bio-Rad) for 30 minutes at room temperature, then incubated overnight at 4°C in blocking solution containing primary antibodies. Membranes were washed three times with TBST (GenDEPOT), followed by incubation with secondary antibodies; goat anti-mouse/rabbit IRDye (LI-COR Biotechnology, Lincoln, NE), or goat anti-mouse/rabbit-HRP (Jackson ImmunoResearch Laboratories), both diluted 1:10,000 in blocking buffer. After final washes with TBST, membranes were either imaged and analyzed using the Odyssey infrared imaging system (LI-COR Biotechnology) or incubated with enhanced chemiluminescence reagent (Pierce) and scanned using Azure 600 Biosystem (Dublin, CA). All western blots shown are representative examples of three or more experiments.

### PHOSPHOINOSITIDE ELISA ASSAYS

This method was previously described for PI(3)P (He et al., 2016). Retinas from 20 wild-type or knockout mice were collected in 8% (v/v) OptiPrep (Sigma) prepared with Ringer’s solution containing PhosSTOP™ and a protease inhibitor cocktail (Sigma). Retinas were gently vortexed for 2 minutes at low speed and centrifuged at 400 × g for 2 minutes at room temperature. The supernatants were collected on ice. This process was repeated five times, followed by layering onto a 10–30% (v/v) OptiPrep gradient and centrifugation for 60 minutes at 19,210 x g at 4 °C using a TLS-55 rotor (Beckman Coulter). The ROS (rod outer segments) were collected, diluted in Ringer’s buffer containing PhosSTOP, and pelleted using the TLS-55 rotor for 30 minutes at 32,172 x g at 4 °C. The pellet was resuspended in 1 ml Ringer’s buffer containing PhosSTOP, mixed with 1 ml chloroform:methanol (1:2, v/v), and vortexed for 2 minutes at room temperature. Samples were centrifuged for 10 minutes at 3,661 × g, and the lower organic phase was transferred to a glass tube. This extraction was repeated three times to remove most phospholipids. Samples were then extracted twice with 1 ml chloroform:methanol:12N HCl (2:4:0.8, v/v/v), vortexed and centrifuged as above. The lower chloroform phase was collected and dried under an argon stream, then stored at −80 °C.

The phosphoinositide ELISA assay in photoreceptor outer segments was performed as previously described (He et al., 2016). Briefly, an inorganic phosphorus assay (Chen et al., 1956) was used to normalize the phosphoinositide extract. 100 pmol of phospholipid, extracted using acidic organic solvents, was dissolved in 40 µl methanol:chloroform (9:1, v/v) and loaded into black PolySorp 96-well plates (Thermo Fisher, 437112). Lipids were dried overnight. The plate was blocked with 5% BSA in PBS for 3 hours at room temperature, then incubated overnight at 4 °C with 100 µl/well of 1 µg/ml purified GST-PLCδ-1D4 in PBS containing 3% BSA. After washing with PBS for 8-10 times, plates were then incubated with 1 µg/ml monoclonal 1D4 antibody in PBS with 3% BSA for 3 hours at room temperature, followed by eight PBS washes. The plates were incubated with 1:3,000 diluted goat anti-mouse HRP-conjugated antibody (Thermo) in PBS with 3% BSA for 1 hour at room temperature. After washing, plates were incubated with 100 µl/well of SuperSignal ELISA Femto Maximum Sensitivity Substrate (Thermo) for 1 minute at room temperature. Chemiluminescence signals were detected using a FlexStation 3 (Molecular Devices). All assays were repeated at least three times.

### DATA ANALYSIS AND STATISTICS

In the phosphoinositide ELISA, the data was normalized with wild-type controls, then data were analyzed using unpaired two-tailed t-tests with GraphPad Prism (v10.6.0) software. For statistical image analysis, integrated fluorescence intensity within a boxed region of interest (ROI) was measured using ImageJ. Intensities from PIP5K knockout images were normalized to control ROIs. Statistical comparisons were performed using Student’s t-test in GraphPad Prism, with p < 0.05 considered significant. Data are presented as mean ± standard deviation.

