## Supplementary figures and images for "Differential and compensatory roles for type I phosphatidylinositol-4-phosphate-5-kinase isoforms in retinal function and health"

### Supplemental Figure S1

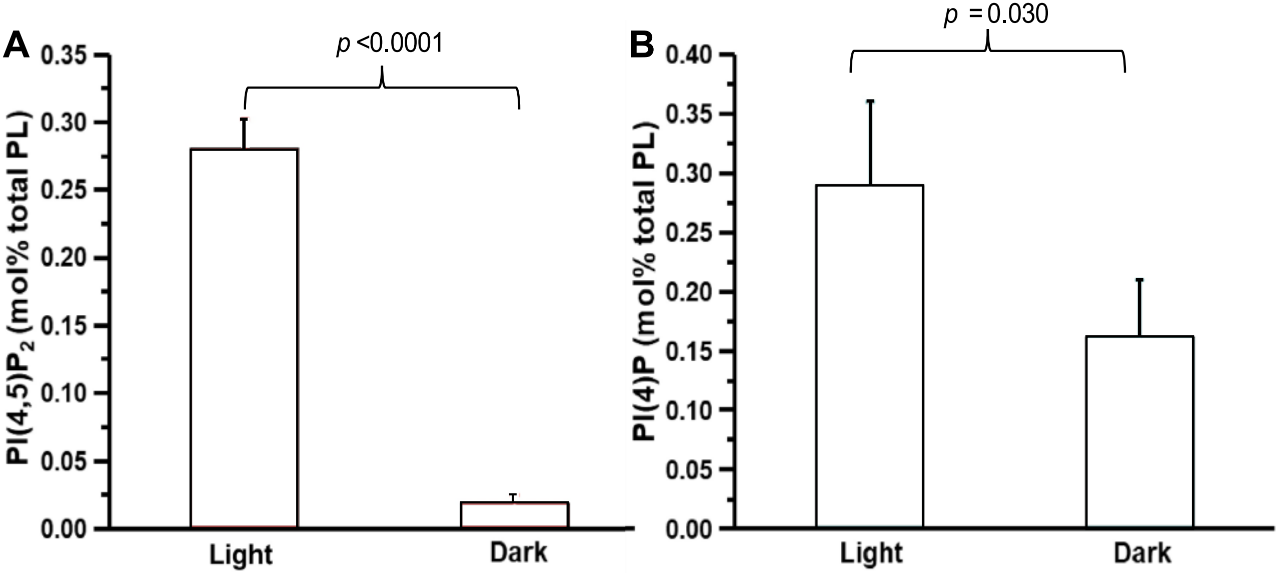

### Supplemental Figure S4

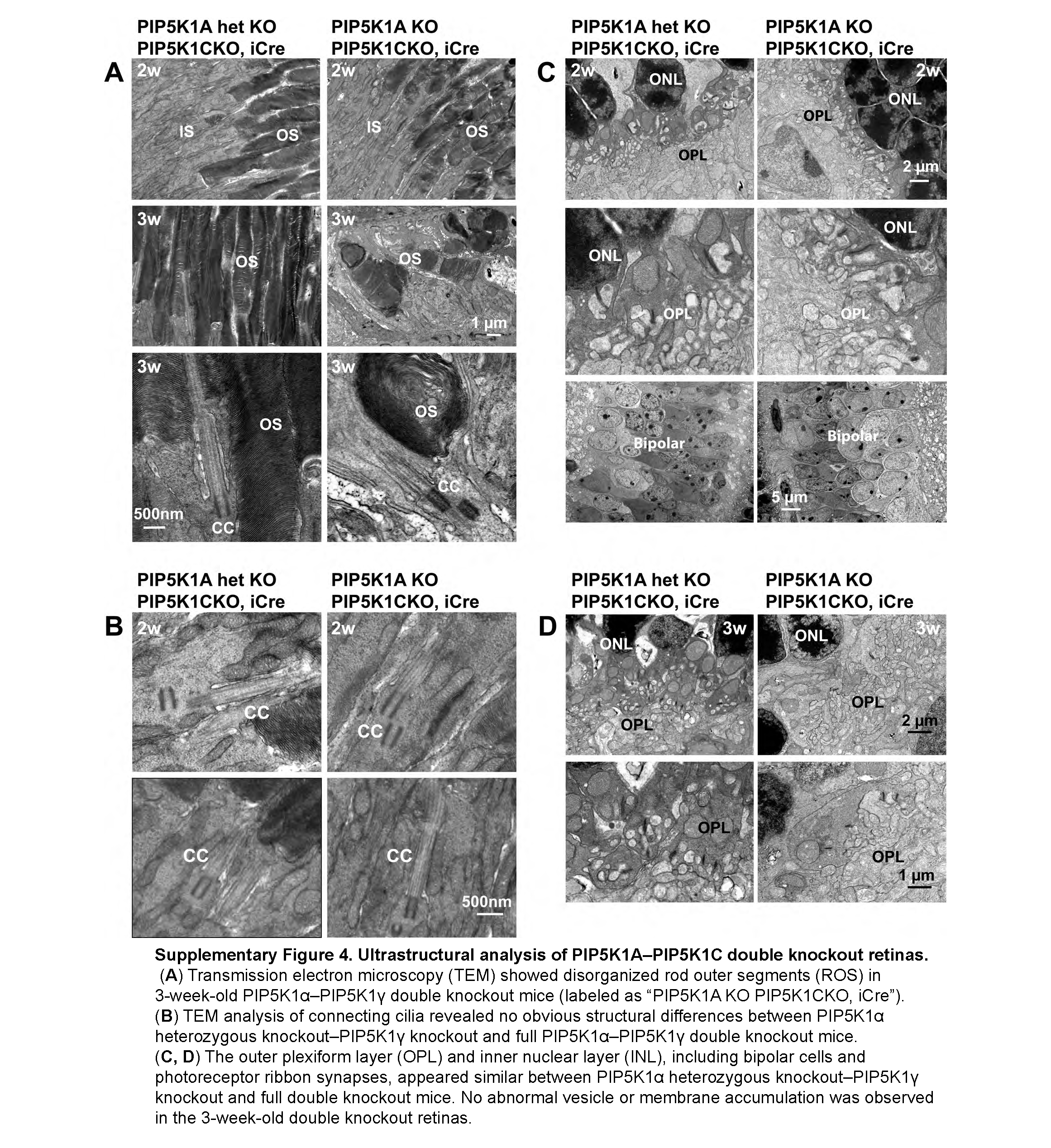

### Supplemental Figure S5

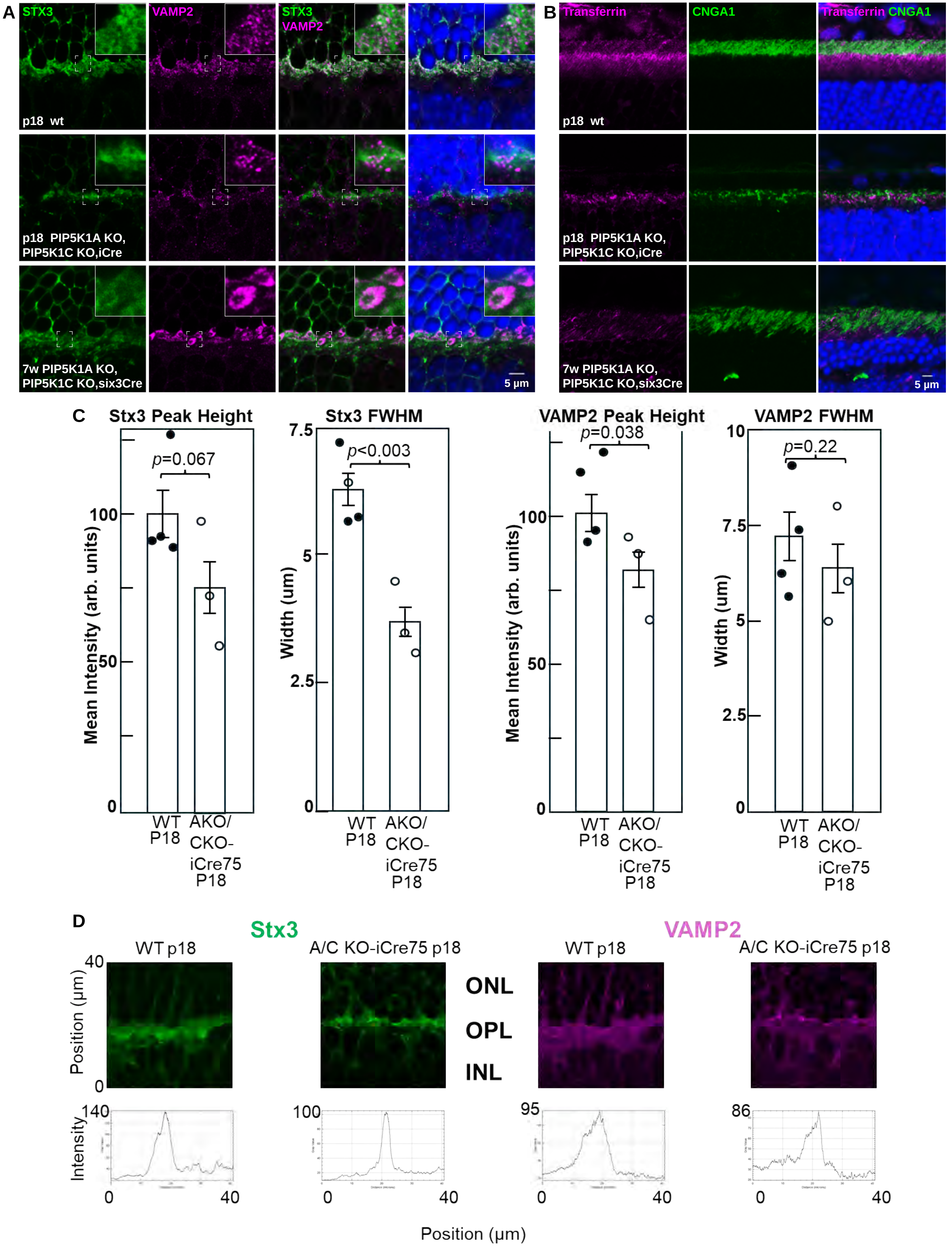

### Supplemental Figure S6

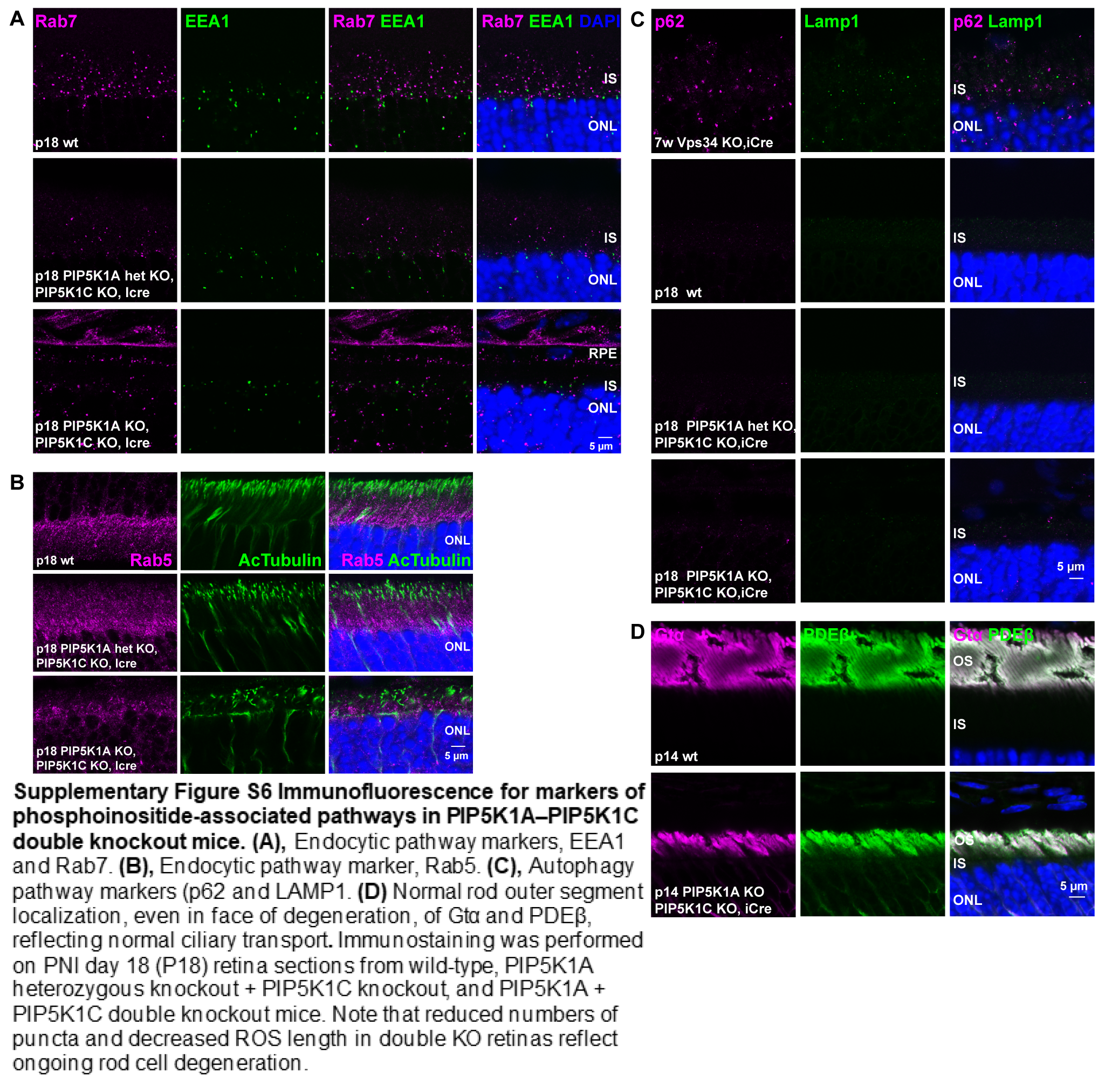

### Supplemental Figure S8

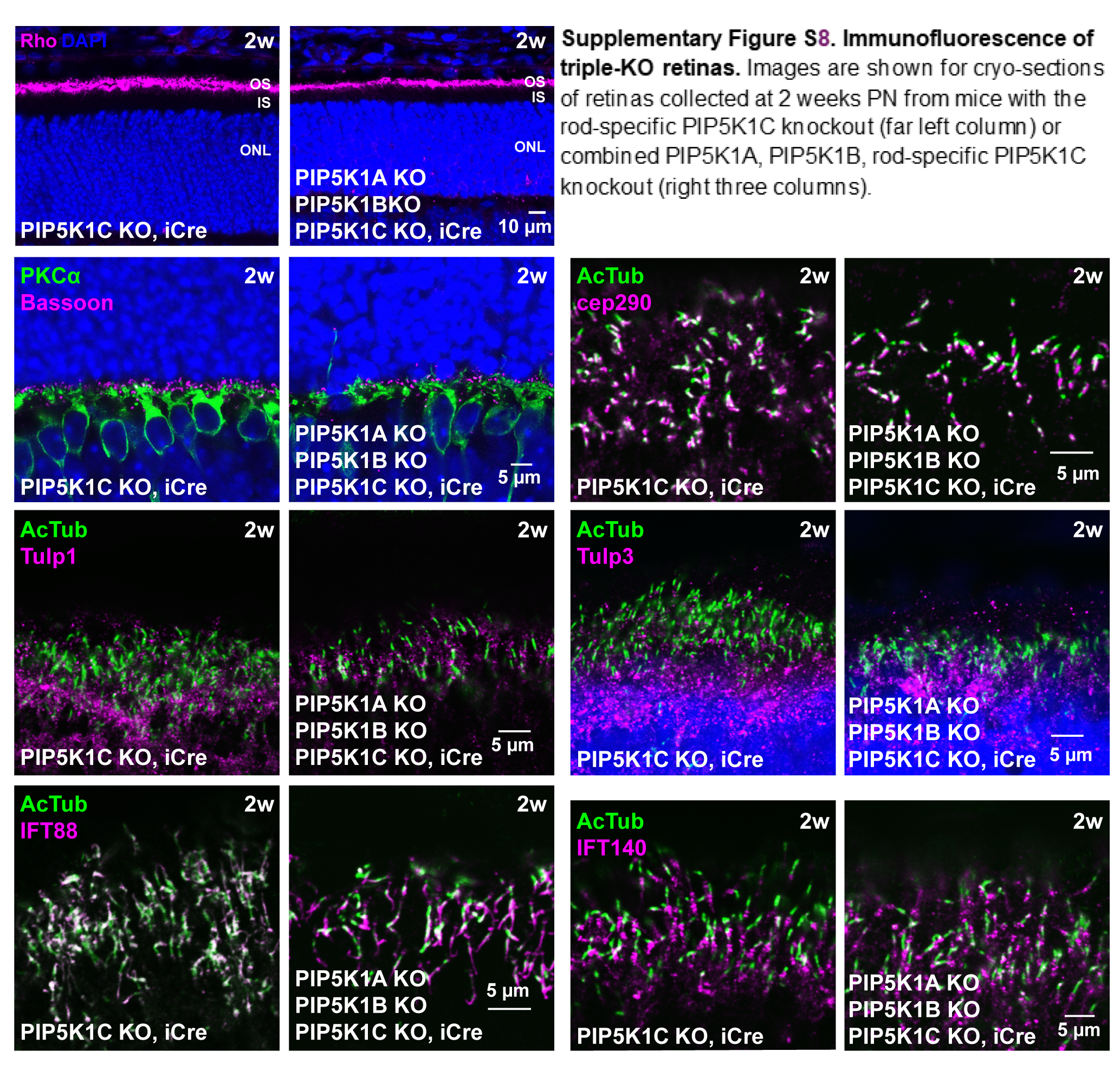
