## Supplemental Figure S2 for "Differential and compensatory roles for type I phosphatidylinositol-4-phosphate-5-kinase isoforms in retinal function and health"

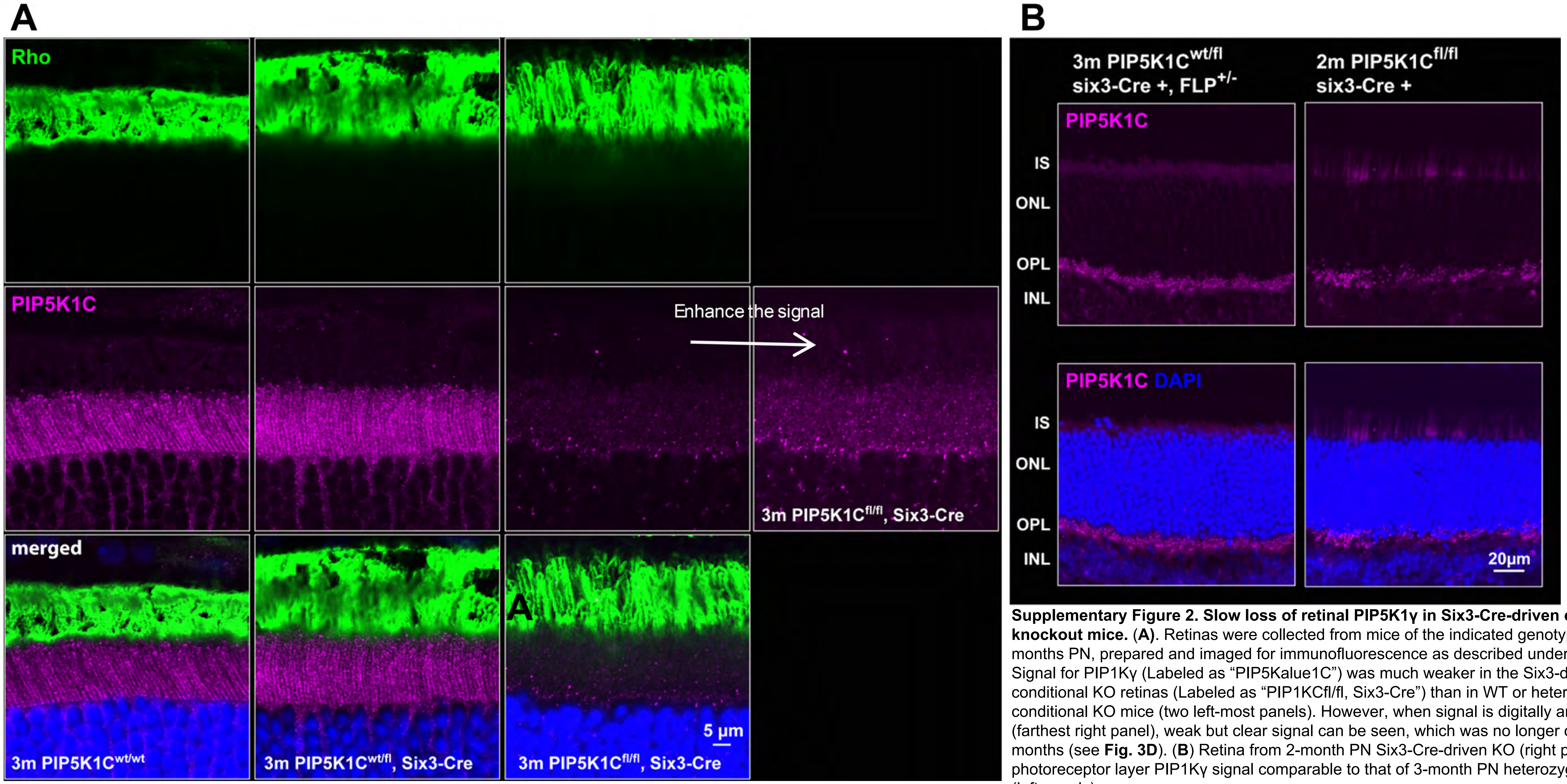


**Supplementary Figure 2. Slow loss of retinal PIP5K1γ in Six3-Cre-driven conditional knockout mice.** (**A**). Retinas were collected from mice of the indicated genotypes at 3 months PN, prepared and imaged for immunofluorescence as described under Methods. Signal for PIP1Kγ (Labeled as "PIP5Kalue1C") was much weaker in the Six3-driven conditional KO retinas (Labeled as "PIP1KCfl/fl, Six3-Cre") than in WT or heterozygous conditional KO mice (two left-most panels). However, when signal is digitally amplified (farthest right panel), weak but clear signal can be seen, which was no longer observed at 5 months (see **Fig. 3D**). (**B**) Retina from 2-month PN Six3-Cre-driven KO (right panels) shows photoreceptor layer PIP1Kγ signal comparable to that of 3-month PN heterozygous controls (left panels).
