## Supplemental Figure S3 for "Differential and compensatory roles for type I phosphatidylinositol-4-phosphate-5-kinase isoforms in retinal function and health"

**Supplementary Figure 3. Regional variation in retinal degeneration in PIP5K1A–PIP5K1C double knockout mice.** DAPI staining of nuclei in the outer nuclear layer (ONL) of PIP5K1α–PIP5K1γ double knockout (labeled as "PIP5K1A KO PIP5K1C KO") photoreceptor cells revealed accelerated retinal degeneration in the dorsal (**A**) and temporal (**B**) regions compared to other retinal areas in 3-week-old double knockout mice. Images of multiple fields were collected to cover the entirety of pan-retinal sections, and digitally "stitched" together to generate the images shown.

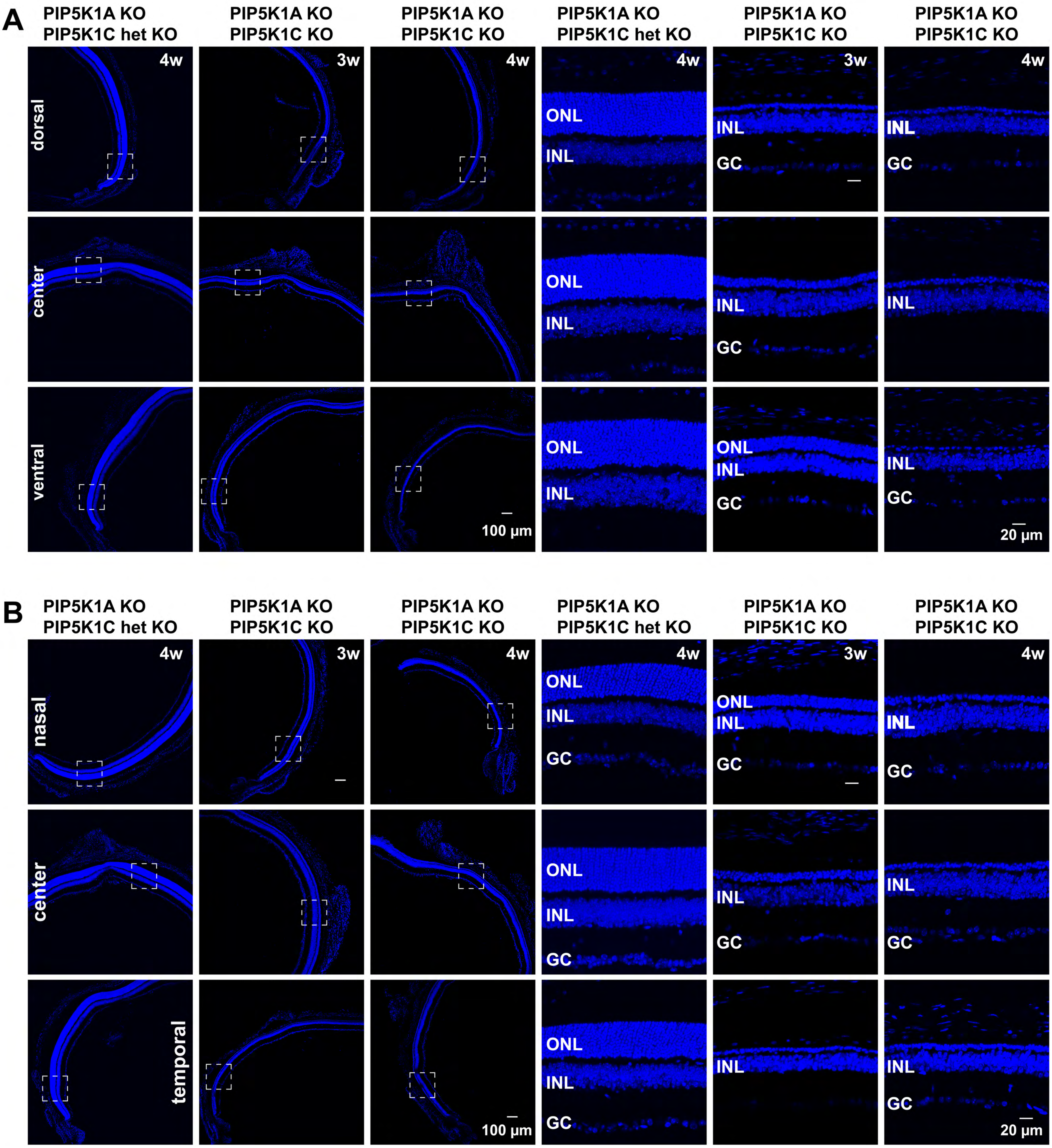
