## Supplemental Figure S7 for "Differential and compensatory roles for type I phosphatidylinositol-4-phosphate-5-kinase isoforms in retinal function and health"

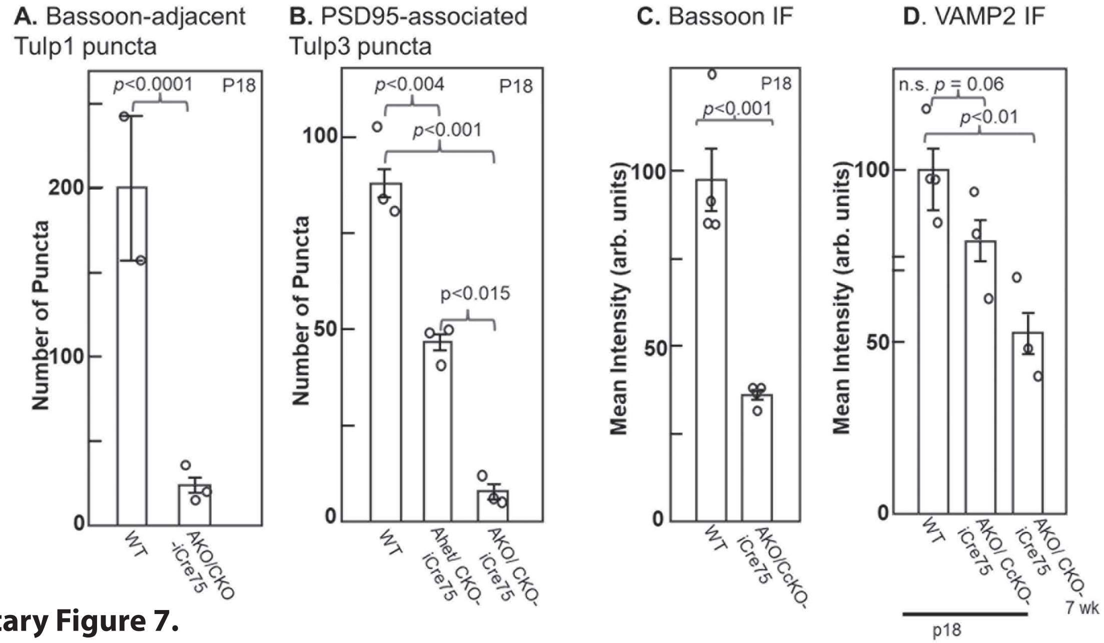


**Supplementary Figure 7. Quantification of alteration in immunofluorescence signals for synaptic markers in the OPL.**

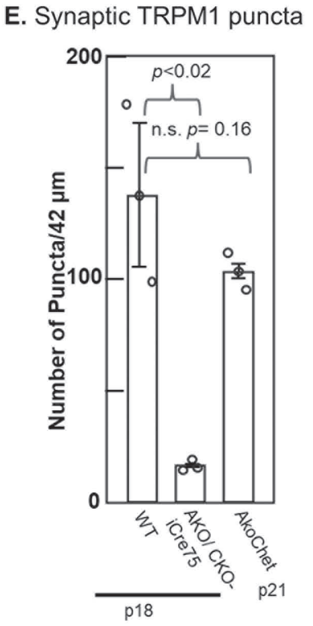


In retinas of the indicated ages and genotypes, immunofluorescence staining and DAPI staining of nuclei were used to identify the synaptic regions of the OPL (**A, B**), staining for Tubby-like PI(4,5)$P_2$-binding proteins. A, Number of Bassoon-adjacent puncta of post-synaptic Tulp3 per 51 µm circumferential length of OPL field. B, PSD-95-associated puncta of pre-synaptic Tulp1 per 42 µm circumferential width of field. (**C**). Integrated intensities for Bassoon immunostaining in 41 µm x 14.6 µm strips of OPL.E. (**D**). Integrated intensities for VAMP2 immunostaining in 41 µm x 14.6 µm strips of OPL. (**E**). Number of synaptic TRPM1 puncta, distinguished from ER-resident TRPM1 by adjacency to Bassoon (PN day 18) or PSD-95 (PN day 21). In **A-C**, all retinas were collected at PN day 18, whereas in **D** and **E**, retinas were examined from both PN day 18 and either PN week 7 (**D**) or PN day 21 (**E**).
